# Structural model of microtubule dynamics inhibition by Kinesin-4 from the crystal structure of KLP-12 –tubulin complex

**DOI:** 10.1101/2022.02.14.480441

**Authors:** Shinya Taguchi, Juri Nakano, Tsuyoshi Imasaki, Tomoki Kita, Yumiko Saijo-Hamano, Naoki Sakai, Hideki Shigematsu, Hiromichi Okuma, Takahiro Shimizu, Eriko Nitta, Satoshi Kikkawa, Satoshi Mizobuchi, Shinsuke Niwa, Ryo Nitta

**Author notes:** These authors contributed equally (Shinya Taguchi, Juri Nakano, Tsuyoshi Imasaki).

## Abstract

Kinesin superfamily proteins are microtubule-based molecular motors driven by the energy of ATP hydrolysis. Among them, the kinesin-4 family is a unique motor that inhibits microtubule dynamics. Although mutations of kinesin-4 cause several diseases, its molecular mechanism is unclear because of the difficulty of visualizing the high-resolution structure of kinesin-4 working at the microtubule plus-end. Here, we report that KLP-12, a *C. elegans* kinesin-4 ortholog of KIF21A and KIF21B, is essential for proper length control of *C. elegans* axons, and its motor domain represses microtubule polymerization *in vitro*. The crystal structure of the KLP-12 motor domain complexed with tubulin, which represents the high-resolution structural snapshot of inhibition state of microtubule-end dynamics, revealed the bending effect of KLP-12 for tubulin. Comparison with the KIF5B-tubulin and KIF2C-tubulin complexes, which represent the elongation and shrinking forms of microtubule ends, respectively, showed the curvature of tubulin introduced by KLP-12 is in between them. Taken together, KLP-12 controls the proper length of axons by modulating the curvature of the microtubule ends to inhibit the microtubule dynamics.

## Introduction

Kinesin superfamily proteins (KIFs) are microtubule-based molecular motors driven by the energy of ATP hydrolysis (Hirokawa et al., 2009a). Most KIFs move along microtubules to transport various cargos, including membranous organelles, protein complexes and mRNAs (Guedes-Dias and Holzbaur, 2019; Hirokawa et al., 2009b). In addition to transporting cargos, some kinesins possess the ability to regulate microtubule dynamics, such as elongation (polymerization), catastrophe or shrinkage (depolymerization), in diverse ways. Kinesin-1, the founding member of the KIFs, changes the conformation of unstable GDP microtubules to growing GTP microtubules (Muto et al., 2005; Shima et al., 2018). Conversely, kinesin-13 destabilizes both the plus and minus ends of microtubules to induce catastrophe or depolymerization (Desai et al., 1999; Ogawa et al., 2004). Kinesin-8 moves processively toward the microtubule plus-end, where it depolymerizes microtubules (Gupta et al., 2006; Niwa et al., 2012; Varga et al., 2006; Wang et al., 2016).

Kinesin-4, another family of kinesins, is known to inhibit microtubule dynamics and is classified into three subfamilies: the KIF4 subfamily (*Mm*KIF4, *Ce*KLP-19, *Dm*KLP-3A, Xklp1), the KIF7 subfamily (*Mm*KIF7, *Mm*KIF27, *Dm*Cos2), and the KIF21 subfamily (*Mm*KIF21A/B, *Dm*KLP-31E, *Ce*KLP-12) (Yue et al., 2018). The KIF21 subfamily has attracted considerable attention because its genetic alterations are linked with several diseases. Point mutations of KIF21A cause congenital fibrosis of extraocular muscle type 1 (CFEOM1) (Yamada et al., 2003). Polymorphisms in the KIF21B gene are associated with several inflammatory diseases, such as multiple sclerosis or Crohn’s disease (Barrett et al., 2008; Goris et al., 2010). Increased expression of KIF21B is also linked to the progression of neurodegenerative disorder (Kreft et al., 2014). *Kif21b* knockout mice were reported to exhibit behavioral changes involving impaired learning and memory (Muhia et al., 2016).

The molecular mechanisms of how kinesin-4 affects microtubule dynamics have been studied for more than a dozen years, demonstrating the main contribution of their motor domains to microtubule dynamics inhibition. Xklp1/KIF4, a fast processive motor, was first reported to reduce both microtubule growth and the catastrophe rate (Bieling et al., 2010; Bringmann et al., 2004). Its motor domain is able to bind not only to the microtubule lattices for microtubule-based motility but also to the curved tubulin dimers for inhibition of microtubule dynamics. Nonmotile KIF7 was reported to reduce the microtubule growth rate but enhance catastrophe to organize the tips of ciliary microtubules (He et al., 2014; Yue et al., 2018). The processive motor KIF21A/B reduces microtubule growth and catastrophes similar to Xklp1/KIF4 (van der Vaart et al., 2013; van Riel et al., 2017). These studies suggest that the motor domains of kinesin-4 family proteins play a crucial role in reducing the growth rate of microtubules. In other words, minor alterations in kinesin-4 motor domains affect their conserved functions to suppress microtubule dynamics by displaying strikingly distinct motility characteristics (van der Vaart et al., 2013; van Riel et al., 2017).

The other domains of kinesin-4 are also known to be involved in microtubule dynamics inhibition by regulating or supporting motor domain functions. The coiled-coil region of KIF21A in which CFEOM1-associated mutations are localized operates as an autoinhibitory domain by interaction with the motor domain (Bianchi et al., 2016; Cheng et al., 2014; van der Vaart et al., 2013). The dominant character of CFEOM1 syndrome is thus connected to the increased activity of the mutant KIF21A kinesin caused by the loss of autoinhibition. The WD40 domain of KIF21B holds on to the growing microtubule tip and induces its pausing (van Riel et al., 2017), which is required for the sustained action of the motor domain on the microtubule plus-end to inhibit microtubule dynamics.

We previously reported the first crystal structure of the KIF4 motor domain (Chang et al., 2013) and showed molecular mechanisms of ATP-induced motion; however, because of the lack of functional and structural information on kinesin-4 on the microtubule plus-end, the mechanism by which kinesin-4 motors inhibit microtubule dynamics is still obscure. Here, we investigated the functional and structural analyses of microtubule dynamics inhibition by kinesin-4 KLP-12, a *Caenorhabditis elegans (C. elegans)* ortholog of KIF21A and KIF21B (Figure 1A; Figure 1–figure supplement 1). Genetic analyses and *in vitro* TIRF (Total Internal Reflection Fluorescence microscopy) assays showed that KLP-12 regulates axonal length through inhibiting microtubule dynamics at its plus-end, similar to KIF21A and KIF21B. The crystal structure of KLP-12 complexed with curved α-, β-tubulin dimers suggested the structural model of microtubule dynamics inhibition by the kinesin-4 motor domain; kinesin-4 precisely controls the curvature of tubulin dimers at the plus-end, which is larger than that decorated by plus-end stabilizing kinesin-1 and smaller than that decorated by destabilizing kinesin-13. This precise control was achieved by the specific interactions on the microtubule interfaces conserved among kinesin-4 motors.

**Figure 1.**
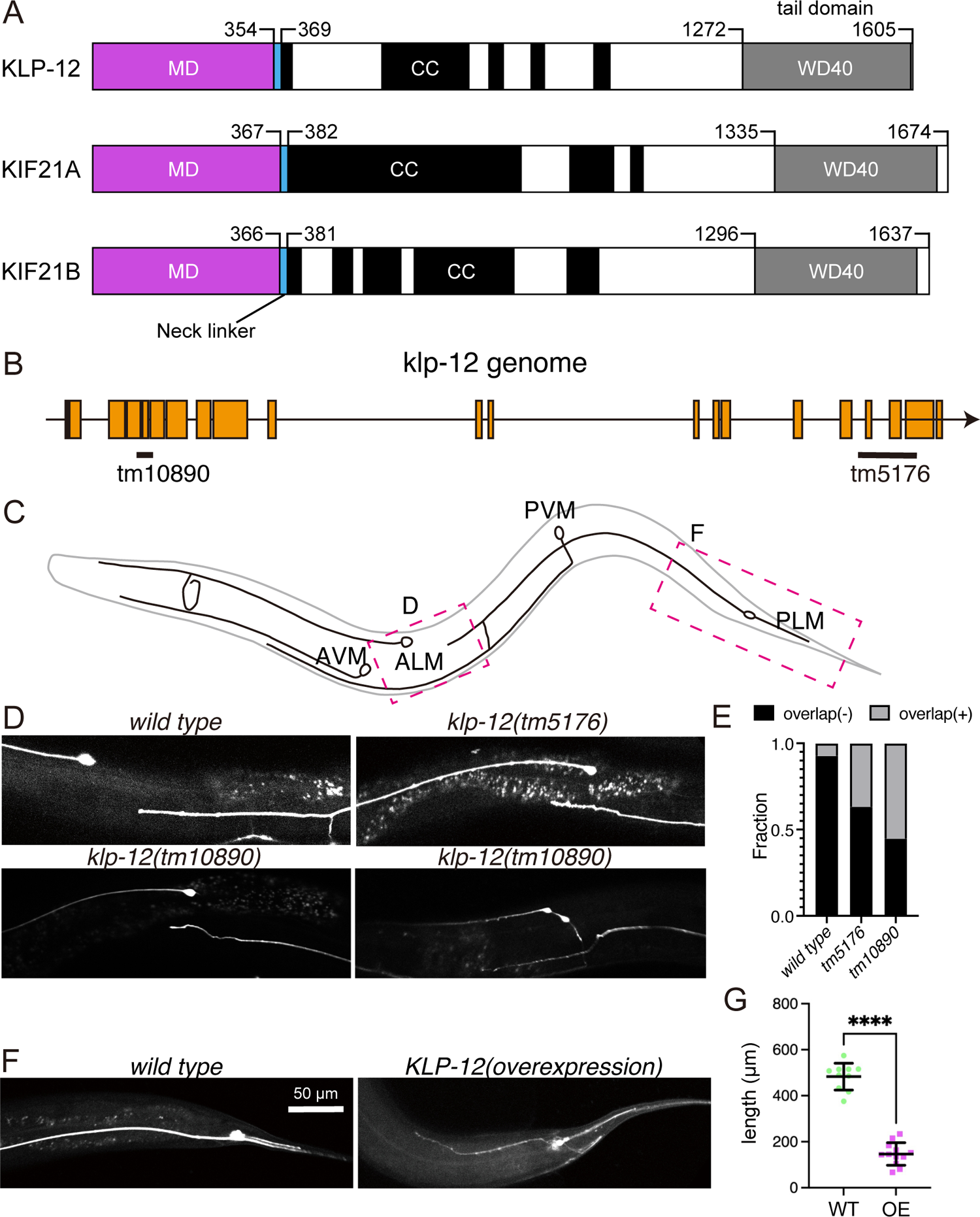
KLP-12 is an ortholog of KIF21A and KIF21B that regulates axonal length. (A) Schematic presentation of the domain organization of the KIF21 subfamily: *C. elegans* KLP-12, human KIF21A, and KIF21B consist of a motor domain (MD; magenta), neck linker (blue), coiled-coil domains (CC; black), and WD40 domain (WD40; gray). Phylogenetic tree and sequence alignment of kinesin-4 family are available in Figure 1 - figure supplement 1 and Figure 1 - figure supplement 2, respectively. (B) Schematic presentation of the genomic structure of *C. elegans klp-12* and mutant alleles used in this study. (C) Schematic presentation of the sensory neurons. Areas observed in panels (D) and (F) are shown by magenta boxes. (D) The tiling of anterior lateral microtubule cell (ALM) and posterior lateral microtubule cell (PLM) neurons. The axonal tip of PLM neurons does not overlap with the cell body of ALM neurons in the wild type, while the axonal tip overlaps with the ALM cell body in *klp-12(tm5176)* and *klp-12(tm10890)*. (E) The percentage of ALM and PLM overlap in wild-type, *klp-12(tm5176),* and *klp-12(tm10890)* mutant worms. (F) Overexpression of KLP-12 suppresses the elongation of the mechanosensory axon. (G) Quantification of the length of neurons in wild-type (WT) and KLP-12-overexpressing (OE) *C. elegans*.

**Figure 1 - figure supplement 1.**
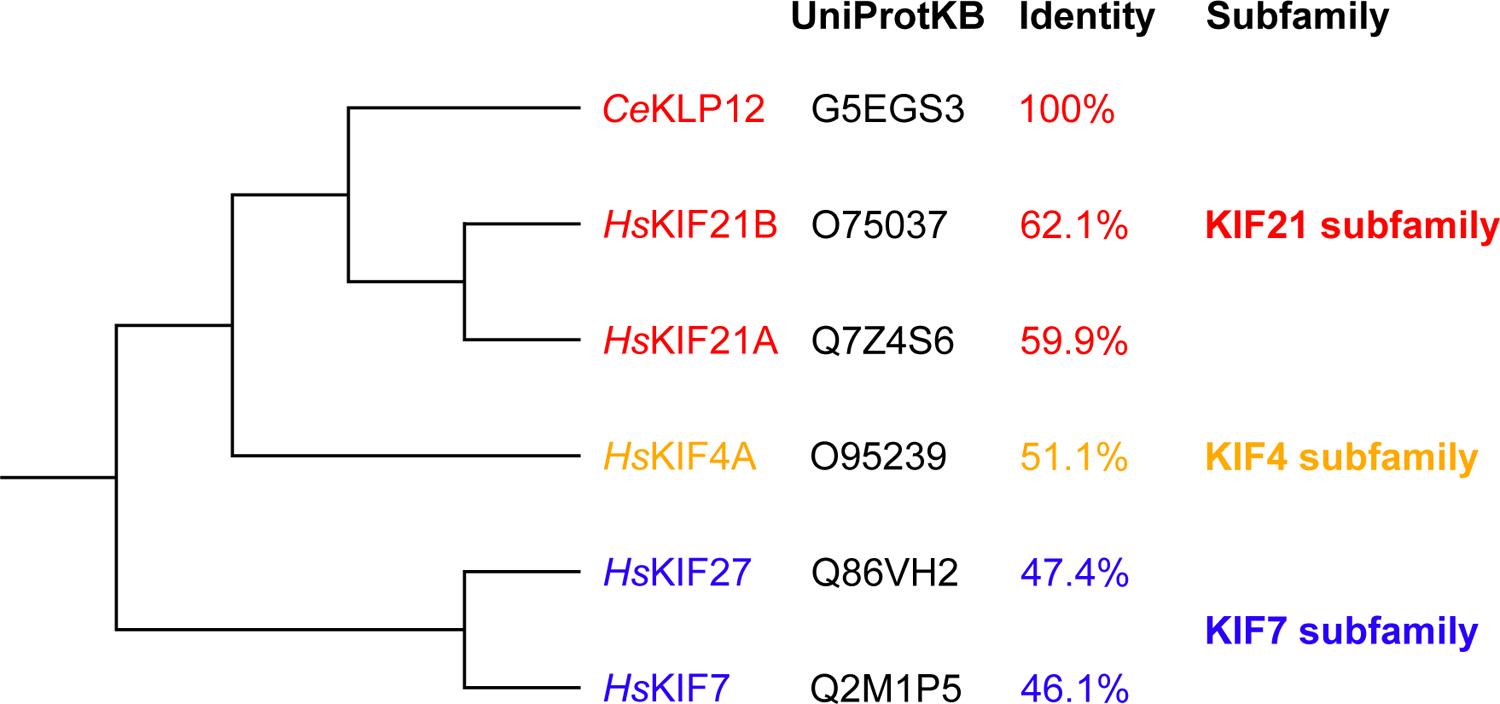
Phylogenetic tree of representative kinesin-4 family member motor domains. This phylogenetic tree was created with reference to Yue et al., 2018. The identities were calculated by SIM (https://web.expasy.org/sim/).

**Figure 1 - figure supplement 2.**
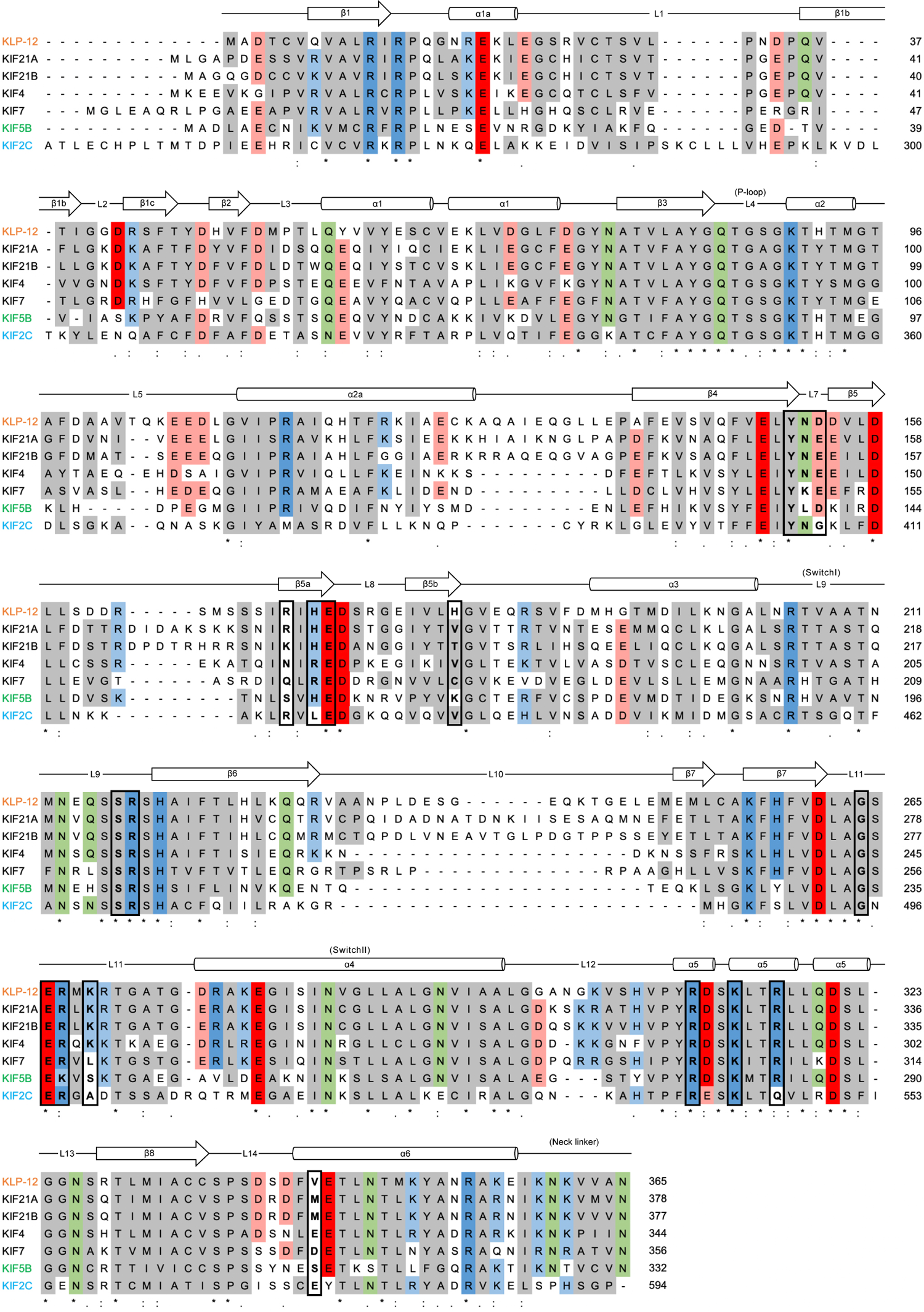
The sequence alignment of kinesin-4 family proteins with KIF5B and KIF2C. The sequence alignment of kinesin-4 family proteins *Ce*KLP-12, *Hs*KIF21A, *Hs*KIF21B, *Mm*KIF4, *Mm*KIF7, Kinesin-1 family *Hs*KIF5B, and kinesin-13 family *Hs*KIF2C motor domain with secondary structure of KLP-12(M). Well conserved residues among Kinesin-4 motors are indicated by coloring. Alignment was generated by Clustal Omega (https://www.ebi.ac.uk/Tools/msa/clustalo/).

## RESULTS

### KLP-12 regulates the length of axons in *C. elegans* neurons

KLP-12 is predicted to be a worm orthologue of KIF21A and KIF21B which regulates axonal length. However, the function of KLP-12 remains to be totally elusive. Thus, we firstly analyzed the phenotype of *klp-12* mutants. We used two independent mutant alleles of *klp-12*, *klp-12(tm10890)* with a defect in the motor domain, and *klp-12(tm5176)* with a defect in the tail domain (Figure 1B). We observed the development of two mechanosensory neurons, anterior lateral mechanosensory (ALM) and posterior lateral mechanosensory (PLM) neurons (Figure 1C). In wild-type animals, the axonal tip of PLM neurons does not reach the cell body of ALM without overlapping with each other (wild-type in Figure 1D and E) (Gallegos and Bargmann, 2004). On the other hand, more than 30% of the PLM axon in *C. elegans* with *klp-12(tm5176)* overtook the cell body of ALM (*klp-12(tm5176)* in Figure 1D and E). Motor domain-deficient *klp-12* (*klp-12(tm10890)*) showed a more severe phenotype; more than 50% of neurons overlapped, and thin warped axons were observed (*klp-12(tm10890)* in Figure 1D and E). The results also suggested that both the motor domain and the tail domain are necessary to precisely control axon length. We also investigated the effect of overexpression of wild-type KLP-12 in *C. elegans* neurons. Compared to the wild type, the PLM axon overexpressing KLP-12 became strikingly shorter and thinner (Figure 1F and G). Taken together with these results, the appropriate activity of KLP-12 is necessary to achieve proper length control of axons, suggesting that the function of KIF21/KLP-12 family proteins (Figure 1A) is evolutionarily conserved.

### KLP-12 is a plus-end directed motor that represses microtubule polymerization

Mammalian orthologs of KLP-12, KIF21A and KIF21B, regulate the axon length by inhibiting the microtubule polymerization (Cheng et al., 2014; van Riel et al., 2017). Thus, KLP-12 may also inhibit the microtubule polymerization to restrict the length of axons. To directly visualize the KLP-12 function on the microtubules, we observed the microtubule-based motility of KLP-12 and measured the growth rate of dynamic microtubules with or without KLP-12 in an *in vitro* assay combined with TIRF microscopy (Figure 2). Since the neck-coiled coil sequence of KLP-12 is not sufficient to form a stable dimer, we intended to introduce the leucine zipper (LZ) of GCN4 after KLP-12 (1-393) the neck-coiled coil for dimerization (Tomishige et al., 2002), which is a similar technique used in the previous kinesin-4 family motor study (Yue et al., 2018).

**Figure 2.**
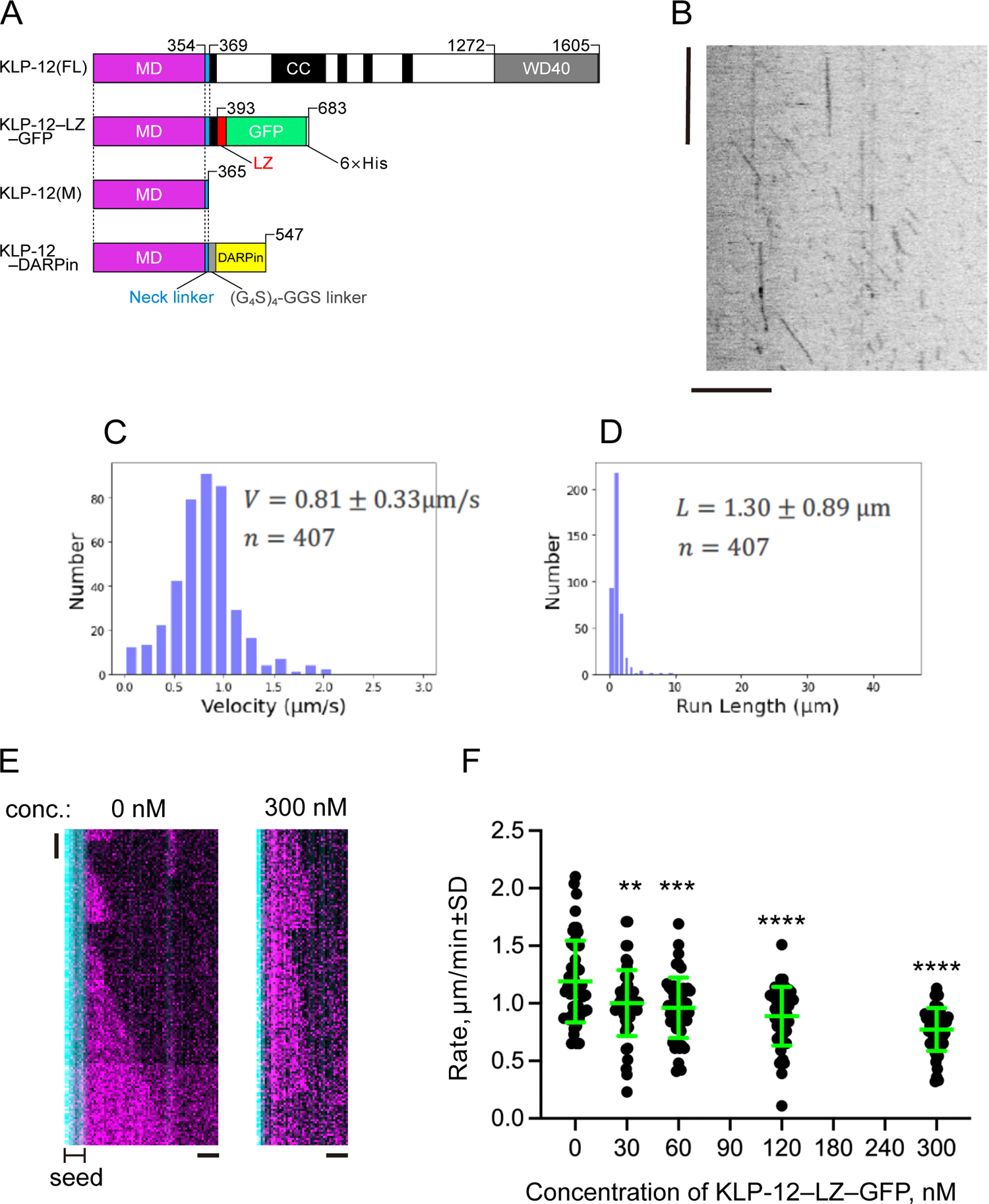
KLP-12 is a plus-end directed motor that represses microtubule polymerization. (A) Schematic presentation of full-length KLP-12 (KLP-12(FL)), KLP-12 (1-393) with GFP connected with a leucine zipper (KLP-12–LZ–GFP), KLP-12 motor domain (KLP-12(M)), and KLP-12 (1-365) with DARPin connected with a flexible linker (KLP-12–DARPin). (B-D) The motility of KLP-12–LZ–GFP on microtubules observed by TIRF microscopy. (B) A representative kymograph showing the motility of KLP-12–LZ–GFP on microtubules. Horizontal and vertical bars respectively show 10 μm and 10 seconds. (C) Histogram showing the velocity of KLP-12–LZ–GFP on microtubules. n = 407 molecules. (D) Histogram showing the run length of KLP-12–LZ–GFP. n = 407 molecules. (E and F) The effect of KLP-12–LZ–GFP on microtubule polymerization. 10 μM of fluorescently labeled microtubules (magenta) were polymerized from GMPCPP stabilized microtubule seeds fixed on the cover glass (cyan). (E) Representative kymographs showing the microtubule polymerization in the presence of 300 nM KLP-12–LZ–GFP (right) or the absence of KLP-12–LZ– GFP (left). Horizontal and vertical bars respectively show 1 μm and 1 minutes. (F) Microtubule growth rate in vitro in the presence of KLP-12–LZ–GFP. Green bars show mean ± standard deviation. **, Adjusted P = 0.0022, ***, Adjusted P = 0.0001, ****, Adjusted P < 0.0001, compared with control (0 nM). One-way ANOVA followed by Dunnett’s multiple comparisons test. n = 52 microtubules. The effect of microtubule growth rate by kinesin-4 family motors is available in Figure 2 - figure supplement 1.

**Figure 2 - figure supplement 1.**
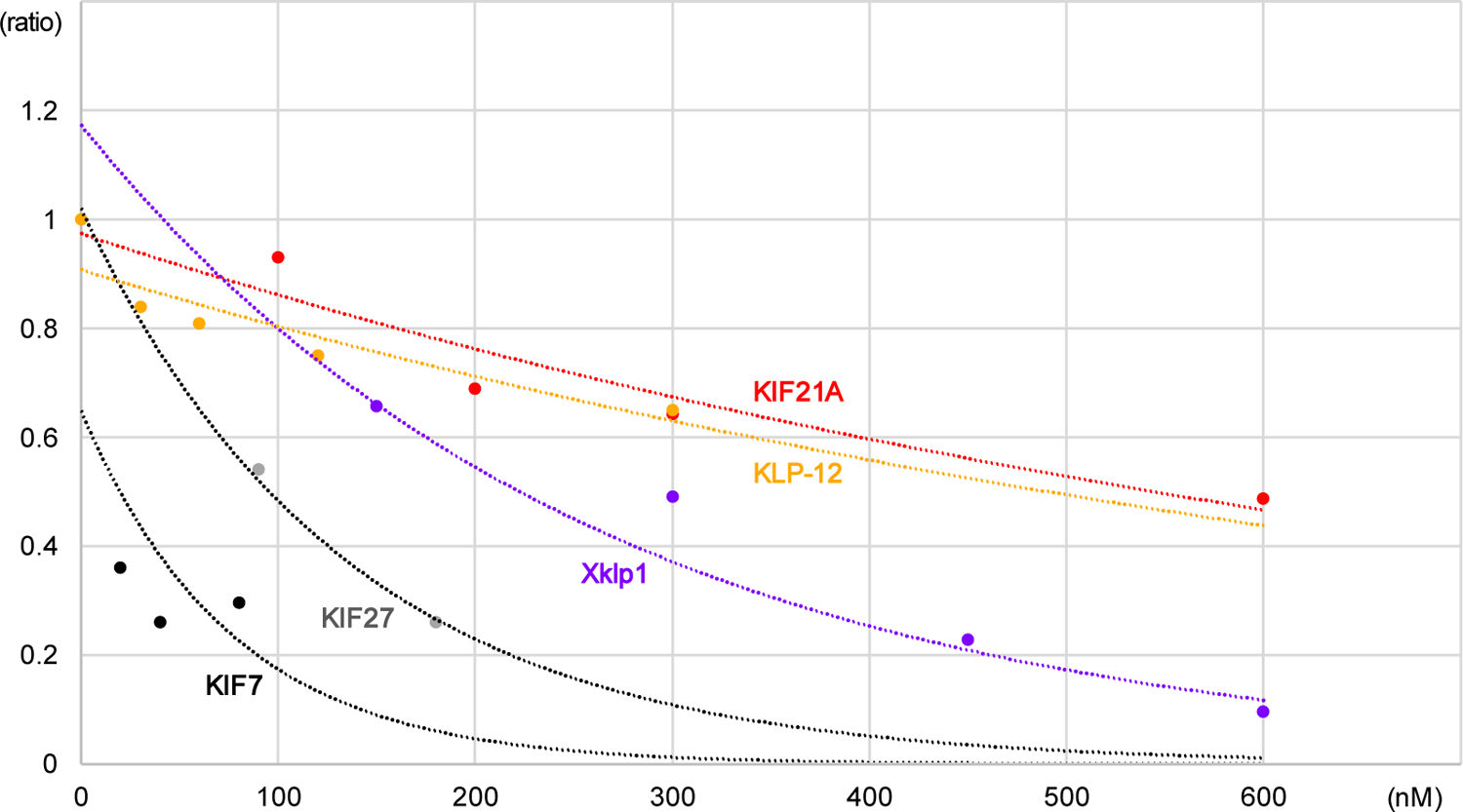
The effect of microtubule growth rate by kinesin-4 family motors.

The LZ and GFP were inserted right after KLP-12 (1-393) (Figure 2A). This KLP-12– LZ–GFP motor showed the plus-end directed movement on the microtubule with the velocity of 0.81 ± 0.33 µm/s and the run length of 1.30 ± 0.89 µm, which is similar speed as the processive fast motor kinesin-1 or KIF4 (Figure 2B-D) (Shima et al., 2018; Yue et al., 2018).

Next, we measured the microtubule dynamics at its plus-end with or without KLP-12– LZ–GFP. In the absence of the KLP-12–LZ–GFP, the microtubule grew at the rate of 1.19 ± 0.35 µm/min (Figure 2E). Increasing amounts of KLP-12–LZ–GFP resulted in inhibiting the microtubule growth rate, with mean rates of 1.00 µm/min at concentrations of 30 nM and 0.77 µm/min at 300 nM, respectively (Figure 2E and F). We further compared the inhibitory effect of KLP-12 with the previously reported dynamics with other kinesin-4 family motors, illustrating that the growth inhibition rate of KLP-12–LZ– GFP was very similar to that of KIF21A (Figure 2 - figure supplement 1), a closely related family with 59.9 % sequence identity to KLP-12 (Figure 1 - figure supplement 1). These results showed KLP-12–LZ–GFP possesses the suppression effect on the microtubule growth rates similar to the other members of the kinesin-4 family proteins, especially to KIF21A/B.

The growth rates of the microtubules were normalized by those of microtubules without kinesin, which were set to the value of 1.0. The vertical axis indicates the ratio of the microtubule growth, and the horizontal axis indicates the kinesin concentration. The current study measures the values of the KLP-12, and the other kinesins were plotted based on the previous studies (Bieling et al., 2010; van der Vaart et al., 2013; Yue et al., 2018).

### The motor domain of KLP-12 binds to both the microtubule lattice and ends to catalyze ATP

Kinesin-4 moves along the microtubule until it reaches the plus-end, at which it inhibits the attachment and release of tubulin-dimers to/from the microtubule end. To achieve these dual functions, kinesin-4 must bind to the microtubule lattice and the microtubule end to catalyze ATP. We thus next focused on the monomeric motor domain of the KLP-12 (KLP-12(M)) (Figure 2A), investigating its ATPase activity in the presence of microtubules or GTP-tubulin dimers. We used GTP-tubulin dimers to investigate the biochemical properties of KLP-12 at the plus-end of microtubules since the plus-end of microtubules is curved due to the lack of lateral interactions between protofilaments, as reported previously (Hunter et al., 2003).

Steady-state ATPase kinetics of KLP-12(M) cells were examined in the presence of GDP microtubules stabilized by paclitaxel (Taxol) (= microtubule lattice) and GTP tubulins (= microtubule end). The basal ATPase activity of KLP-12(M) in the absence of tubulins or microtubules was 0.00023 ± 0.00015 sec^-1^, considerably slower than that of the other kinesin motors (Hunter et al., 2003; Wang et al., 2016). KLP-12(M) ATPase was then activated ∼700 times by microtubules to reach a maximum rate of 0.16 sec^-1^, while that activated more than 1,000 times by free tubulin dimers to reach a maximum rate of 0.29 sec^-1^. *K_M,microtubules_* and *K_M,tubulin_* are 2.7 μM and 8.6 μM (Figure 3A). These results indicate that KLP-12(M) binds similarly to the microtubules and the tubulin-dimers and catalyzes ATP. Thus, KLP-12 could bind both to the microtubule lattice and the microtubule plus-end to activate its ATPase to achieve its dual roles, microtubule motility and stabilization.

**Figure 3.**
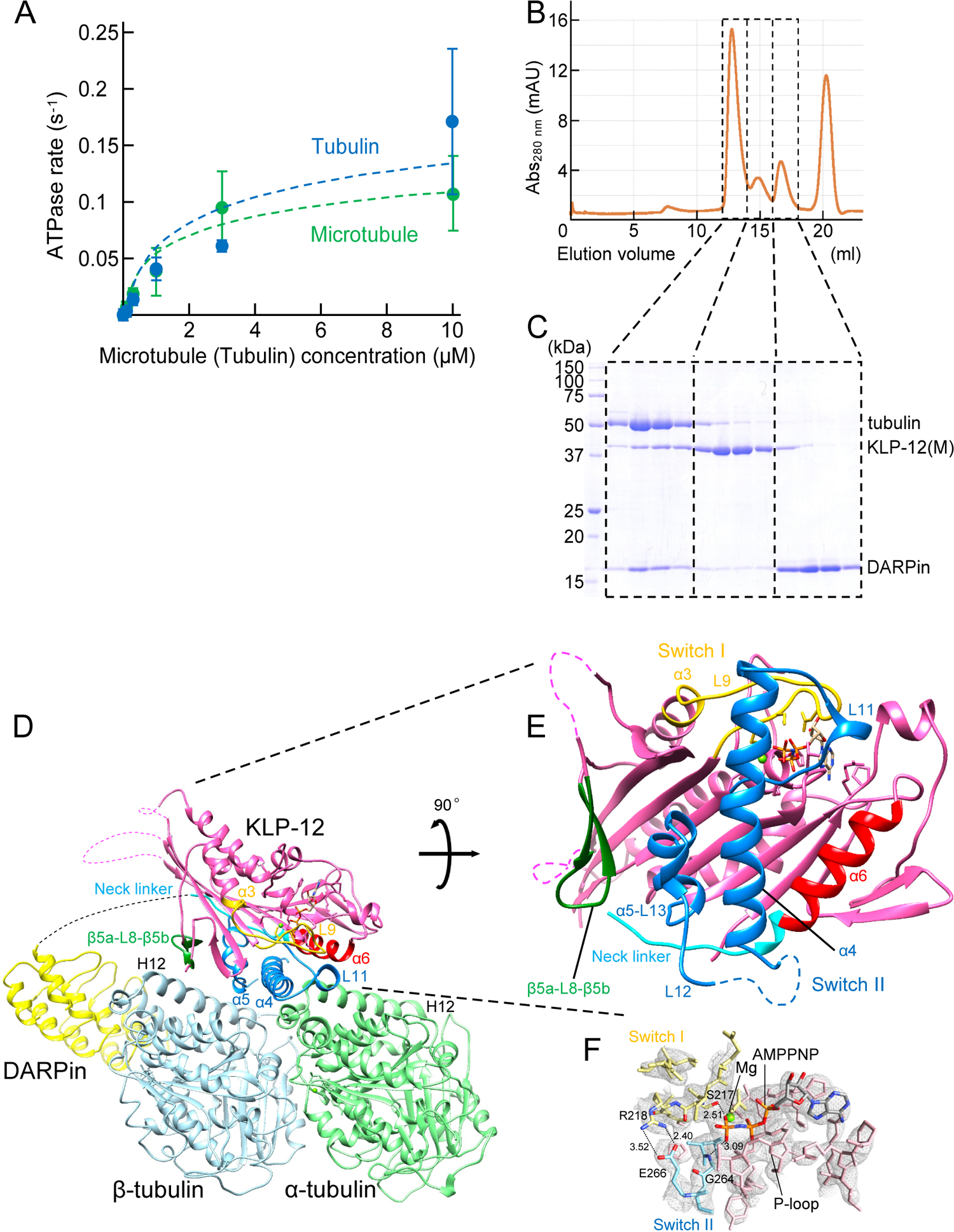
Characterization of KLP-12–DARPin constructs. (A) The steady-state ATPase activity of KLP-12(M) was measured with a microtubule or tubulin heterodimer at 30 °C. Error bars represent standard deviation. Tubulin or microtubule GTPase effect was canceled through the subtracting control without KLP-12(M). (B) Size exclusion chromatography (SEC) of tubulin mixed with KLP-12(M) and DARPin. (C) SDS–PAGE analysis of the SEC peaks of tubulin mixed with KLP-12(M) and DARPin. SDS-PAGE analysis of the SEC peaks of tubulin mixed with KLP-12–DARPin is available in Figure 3 - figure supplement 1. (D) Crystal structure of the tubulin–KLP-12–DARPin complex. Disordered linkers were drawn as a dotted line. α-tubulin is colored light green, β-tubulin is light blue, DARPin is yellow, and KLP-12 is magenta. Structure comparison with KIF4 is available in Figure 3 - figure supplement 2, with KIF5B and KIF2C are available in Figure 3 - figure supplement 3. (E) KLP-12 structure viewed from the interface between tubulins. The residues of the important structure are shown in color. β5a-L8-β5b is green, switch I is yellow, switch II is blue, α6 is red, and the neck linker is cyan. (F) Nucleotide binding pocket of KLP-12. The 2Fo-Fc map around AMP-PNP was calculated with coefficient 2Fo− Fc and contoured at 2.0 σ.

**Figure 3 - figure supplement 1.**
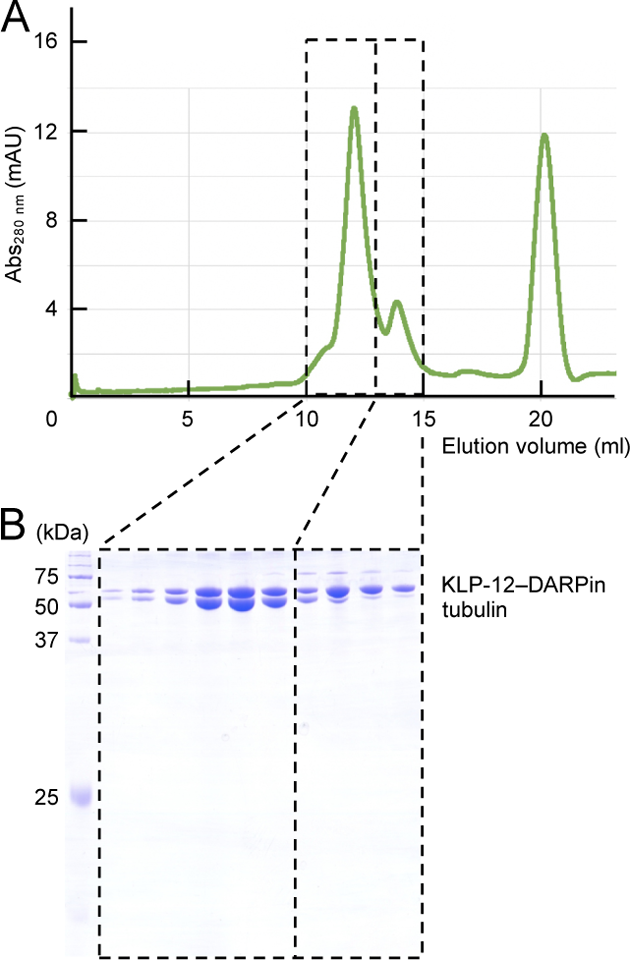
The complex formation of KLP-12–DARPin and tubulin dimers. **A.** Size exclusion chromatography (SEC) of tubulin mixed with KLP-12–DARPin. **B.** SDS-PAGE analysis of the SEC peaks.

**Figure 3 - figure supplement 2.**
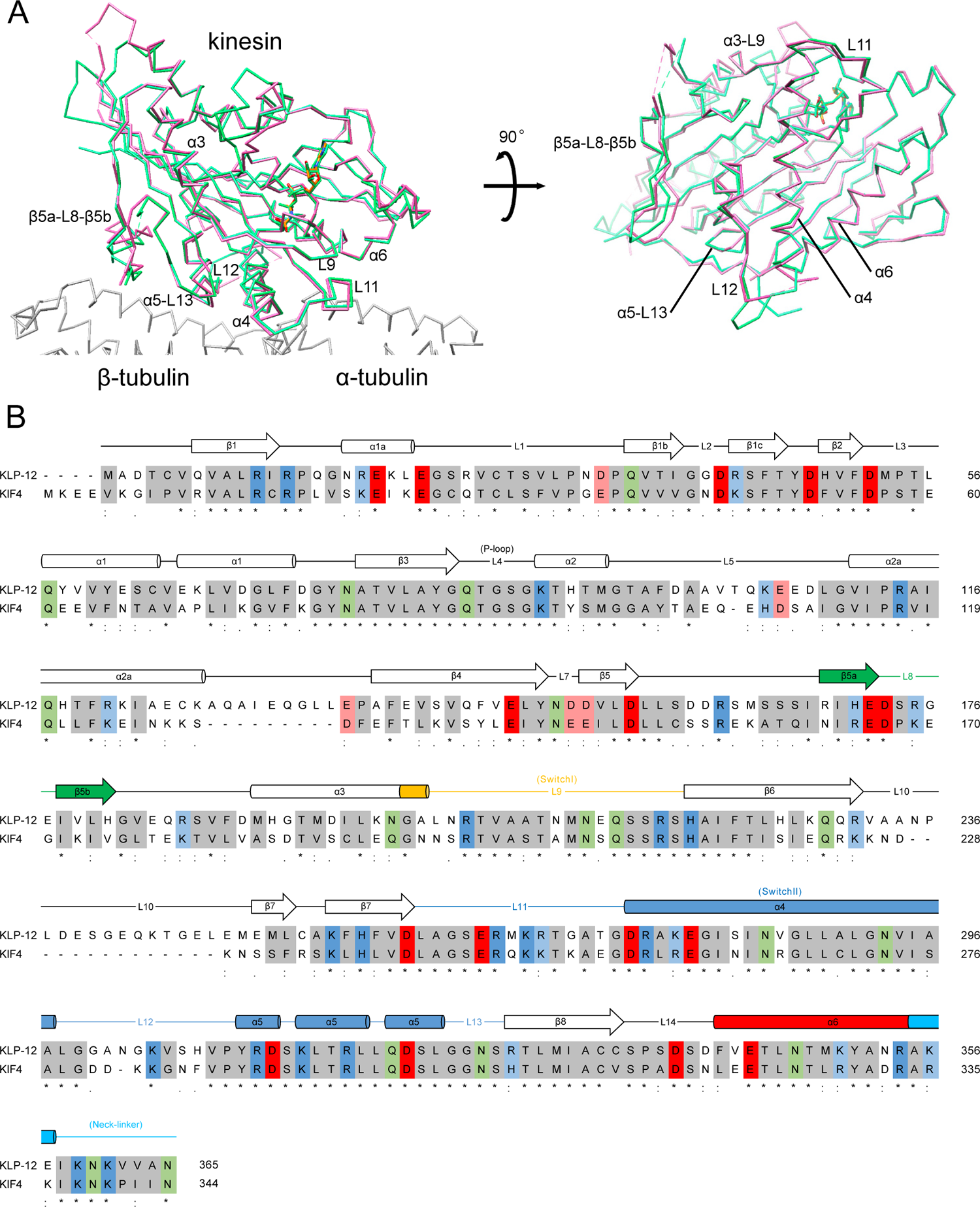
Comparison of motor domains between KLP-12 and KIF4. **A.** Cα chain trace models of *Ce*KLP-12 and *Mm*KIF4 (PDB ID: 3ZFC) motor domain superimposed at kinesin. The average root mean square deviation (RMSD) was 0.974 Å. **B.** The sequence alignment of *Ce*KLP-12 and *Mm*KIF4 motor domain, with structural secondary structure of KLP-12(M). Alignment was generated by Clustal Omega (https://www.ebi.ac.uk/Tools/msa/clustalo/). The important structures are shown in color. β5a-L8-β5b is green, switch I is yellow, switch II is blue, α6 is red, and neck linker is cyan.

**Figure 3 - figure supplement 3.**
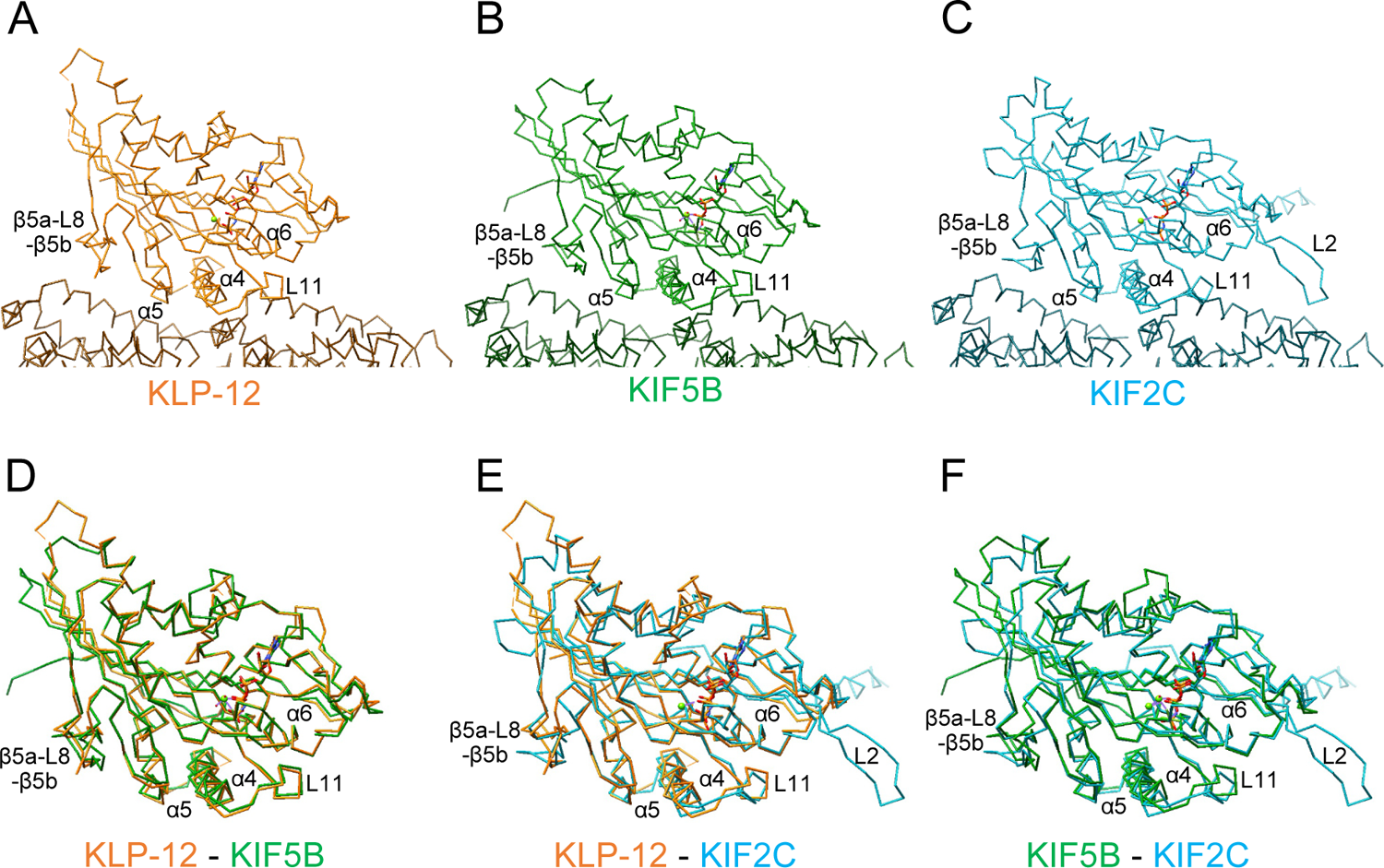
Structure comparison of KLP-12, KIF5B, and KIF2C in complex with tubulin dimers. **A-C.** Cα models of (A) KLP-12, (B) KIF5B, and (C) KIF2C complexes. The KLP-12 complex is in orange, the KIF5B complex (PDB ID: 4HNA) is in green, and the KIF2C complex (PDB ID: 5MIO) is in cyan. **D.** Superimposition of KLP-12 and KIF5B. Root mean square deviation (RMSD) is 1.003 Å. **E.** Superimposition of KLP-12 and KIF2C. RMSD is 1.283 Å. **F.** Superimposition of KIF5B and KIF2C. RMSD is 1.250 Å.

### Structure determination of the tubulin–KLP-12–DARPin complex

The binding ability of the KLP-12 motor domain, KLP-12(M) (Figure 2A), to the soluble tubulin dimer was further analyzed by size exclusion chromatography (SEC). To prevent the self-assembly of tubulins, an A-C2 designed ankyrin repeat protein (DARPin) that binds to the longitudinal interface of β-tubulin was used (Ahmad et al., 2016). Equimolar tubulin dimers, KLP-12(M) with the ATP analog AMP-PNP (adenylyl-imidodiphosphate), and DARPin were injected and analyzed by SEC, representing three peaks (Figure 3B). The first peak consisted of all components, tubulin, KLP-12(M), and DARPin (Figure 3C). The second and third peaks correspond to KLP-12(M) and DARPin, respectively. KLP-12(M) and DARPin apparently shifted to the left side to form a triple complex with tubulin. Thus, KLP-12(M) has a binding ability to a soluble tubulin dimer, consistent with steady-state ATPase assays.

Next, we proceeded to crystallize the KLP-12–tubulin complex to elucidate the molecular mechanism of microtubule stabilization by kinesin-4. For crystallization, to stabilize complex formation between KLP-12(M) and tubulin, KLP-12(M) was fused to DARPin by a long linker. According to a previous study (Wang et al., 2017), the linker length between the C-terminus of KLP-12(M) and the N-terminus of DARPin was optimized.

Four G_4_S repeats with one G_2_S (KLP-12–DARPin fusion construct) were used to form the tubulin–KLP-12–DARPin complex (Figure 2A). Finally, we formed a stable 1:1 complex of KLP-12–DARPin and crystallized tubulin dimers for structure determination (Figure 3–figure supplement 1).

### Crystal structure of Tubulin–KLP-12–DARPin complex

We solved the crystal structure of the tubulin–KLP-12–DARPin complex at 2.9 Å resolution (Figure 3D and E; Table 1). KLP-12 forms an ATP conformation in which switch II takes the up conformation and the neck linker docks to the motor core. The linker between KLP-12 and DARPin, which follows the neck linker, was not observed because of its intrinsic flexibility. In the nucleotide-binding pocket of KLP-12, the density corresponding to AMP-PNP was clearly found with the Mg^2+^ ion (Figure 3F). The highly conserved Ser217 of switch I and Gly264 of switch II are both coordinated to the γ-phosphate of AMP-PNP. The back door between switch I Arg218 and switch II Glu266 was also closed, representing the pre-hydrolysis state during ATP hydrolysis.

**Table 1.** Data collection and refinement statistics

| Tubulin–KLP-12–DARPin |  |  |
| --- | --- | --- |
| Beam line |  | SPRING-8 BL32XU |
| Data collection |  |  |
| Space group | | $P 2_1$ |
| Cell dimensions | $a, b, c$ (Å) | 82.33, 80.98, 117.75 |
| | $\alpha, \beta, \gamma$ (°) | 90.00, 93.51, 90.00 |
| Resolution (Å) |  | 50–2.88 (3.05–2.88) * |
| $R_{\text{meas}}$ (%) | | 35.5 (197.0) |
| $I / \sigma I$ | | 19.28 (3.39) |
| $CC_{1/2}$ | | 97.7 (51.3) |
| Completeness (%) |  | 98.2 (98.8) |
| Redundancy |  | 87.1 (88.7) |
| Refinement |  |  |
| Resolution (Å) |  | 49.25–2.88 |
| No. reflections |  | 33796 |
| $R_{\text{work}}/R_{\text{free}}$ | | 0.211/0.296 |
| No. atoms | Protein | 10301 |
|  | Ligand/ion | 93 |
|  | Water | 0 |
| $B$-factors | Protein | 54.67 |
|  | Ligand/ion | 42.97 |
|  | Water | 0 |
| R.m.s. deviations | Bond lengths (Å) | 0.010 |
|  | Bond angles (°) | 1.40 |

\*Values in parentheses are for the highest-resolution shell.

*C. elegans* KLP-12(M) and the reported structure of *Mus musculus* KIF4 (hereafter called KIF4) complexed with AMP-PNP (PDB ID: 3ZFC) (Chang et al., 2013) were superimposed and compared. The overall conformations of KLP-12 and KIF4 are very similar, with an average root mean square deviation (RMSD) of 0.974 Å (evaluating superpositions across all 319 fully populated columns in the final alignment except disordered residues in the crystal) (Figure 3–figure supplement 2). The microtubule-binding interfaces, which are composed of loop β5a-L8-β5b, switch II (L11-α4-L12-α5-L13), and helix α6, have very similar conformations, and the amino acids in these regions are highly conserved between KLP-12 and KIF4 (Figure 3–figure supplement 2B), reflecting the conserved mechanisms of the microtubule-stabilizing effect of kinesin-4 motors (Bringmann et al., 2004).

### Structural comparisons among three types of kinesin motors, kinesin-4, kinesin-1, and kinesin-13

To elucidate the structural characteristics specific for microtubule-stabilizing kinesin motor kinesin-4, we compared the overall conformations of KLP-12 with the other two types of kinesin motors, kinesin-1 (KIF5B; PDB ID: 4HNA) and kinesin-13 (KIF2C; PDB ID: 5MIO), all of which take the ATP conformation. Kinesin-1 is known to bind preferentially to growing GTP microtubules or stabilize the GDP microtubule (Muto et al., 2005; Peet et al., 2018; Shima et al., 2018), whereas kinesin-13 is known to destabilize microtubules (Ogawa et al., 2004).

Kinesin-4, including KLP-12 except for KIF7, belongs to the plus-end directed processive motors (Yue et al., 2018). The structural comparison showed that the KLP-12 structure resembles plus-end directed kinesin-1 rather than microtubule destabilizing kinesin-13 (Figure 3–figure supplement 3). The RMSDs between KLP-12 and KIF5B (Figure 3–figure supplement 3D) and between KLP-12 and KIF2C (Figure 3–figure supplement 3E) are 1.003 Å and 1.283 Å, respectively, supporting that the binding mode of KLP-12 is slightly more similar to that of KIF5B than that of KIF2C. The low RMSD of KLP-12 and KIF5B indicates that the functional difference in microtubule dynamics is due to the different characteristics of the side chains at the interfaces between kinesin and microtubules. On the other hand, the binding interface of KIF2C is considerably different from KLP-12 and KIF5B because of insertion of loop L2, which is a unique structure of KIF2C (Figure 3–figure supplement 3E and F), serving as an additional binding site to the microtubule (described in a later section).

### Microtubule binding interface of kinesin-4 KLP-12 at α-tubulin

We focused on the interface between α-tubulin and KLP-12 and compared it with the KIF5B and KIF2C structures by superimposing them using the kinesin motor domain (Figure 4; Figure 4–figure supplement 1). KLP-12 Arg267 interacts with tubulin Glu414/Glu420 in H12 with the hydrophobic support of KLP-12 Val343 in α6 through the hydrophobic stem of α-tubulin Glu420 of α-tubulin, albeit Val343 is not conserved well among kinesin-4 (Figure 4A and B). The other interface K269 in L11 which is conserved among KIF21 subfamily and KIF4 subfamily makes a salt-bridge with Glu155 in H4 of α-tubulin. Thus, the KLP-12 structure generates the triangle contacts among L11 of KLP-12, H4 of α-tubulin, and H12 of α-tubulin. The helix α6 of KLP-12 also supports this interface.

**Figure 4.**
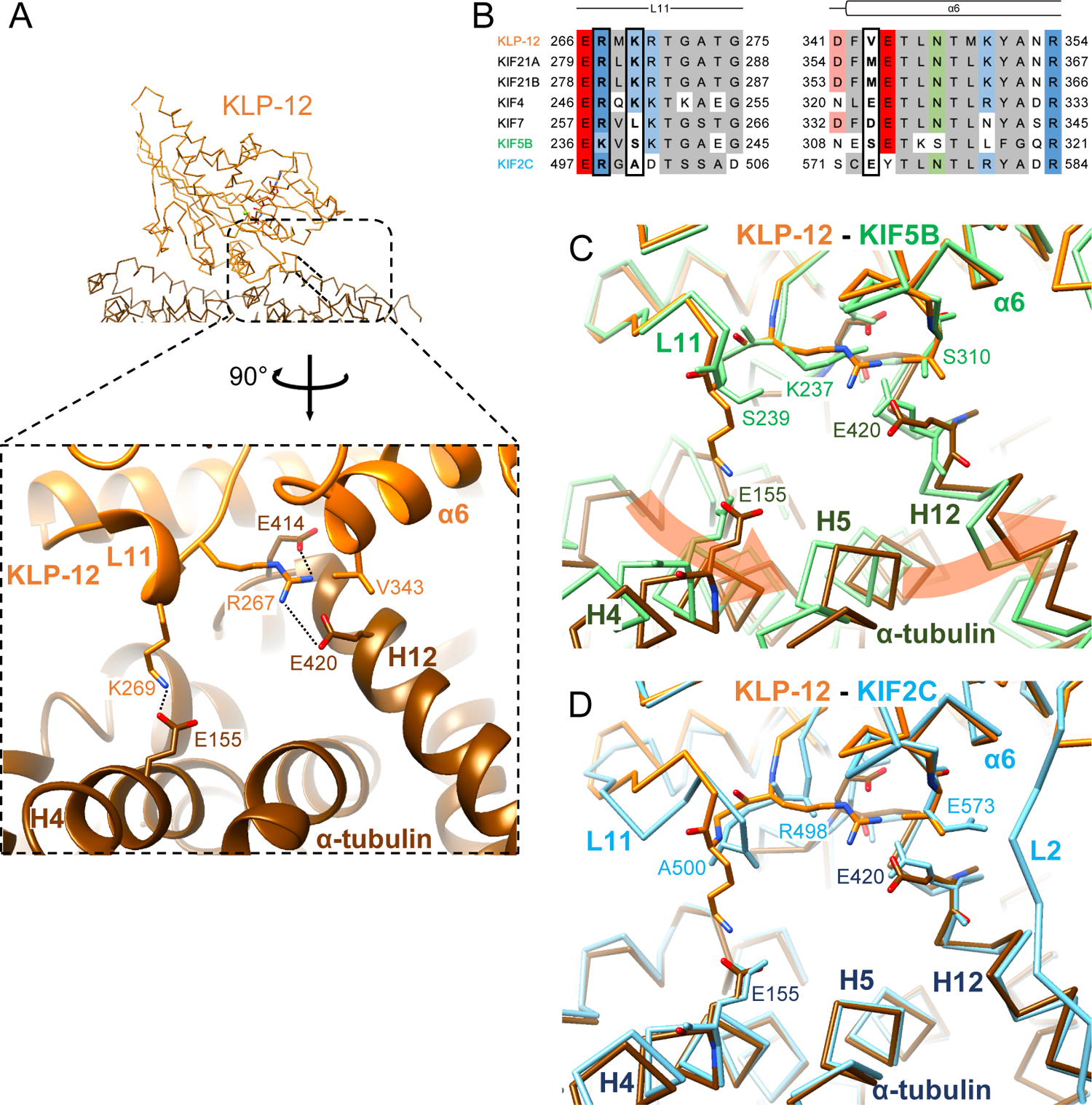
Microtubule binding interface of kinesin-4 KLP-12 at α-tubulin. (A) KLP-12 Arg267 and Lys269 in loop L11 interact with α-tubulin Glu414 and Glu420 in helix H12 and Glu155 in helix H4, respectively. The interfaces of KIF5B and KIF2C at α-tubulin are available in Figure 4 - figure supplement 1. (B) Sequence alignment of the kinesin-4, KIF5B, and KIF2C residues at the interacting area. Interacting residues are marked by squares. (C) Superimposition of Cα chain trace models of the KLP-12 complex (orange) and KIF5B complex (green) at kinesin. The rotation direction of α-tubulin between KLP-12 and KIF5B at helices H4, H5, and H12 is illustrated as a red arrow. (D) Superimposition of Cα chain trace models of the KLP-12 complex (orange) and KIF2C complex (cyan) at kinesin.

**Figure 4 - figure supplement 1.**
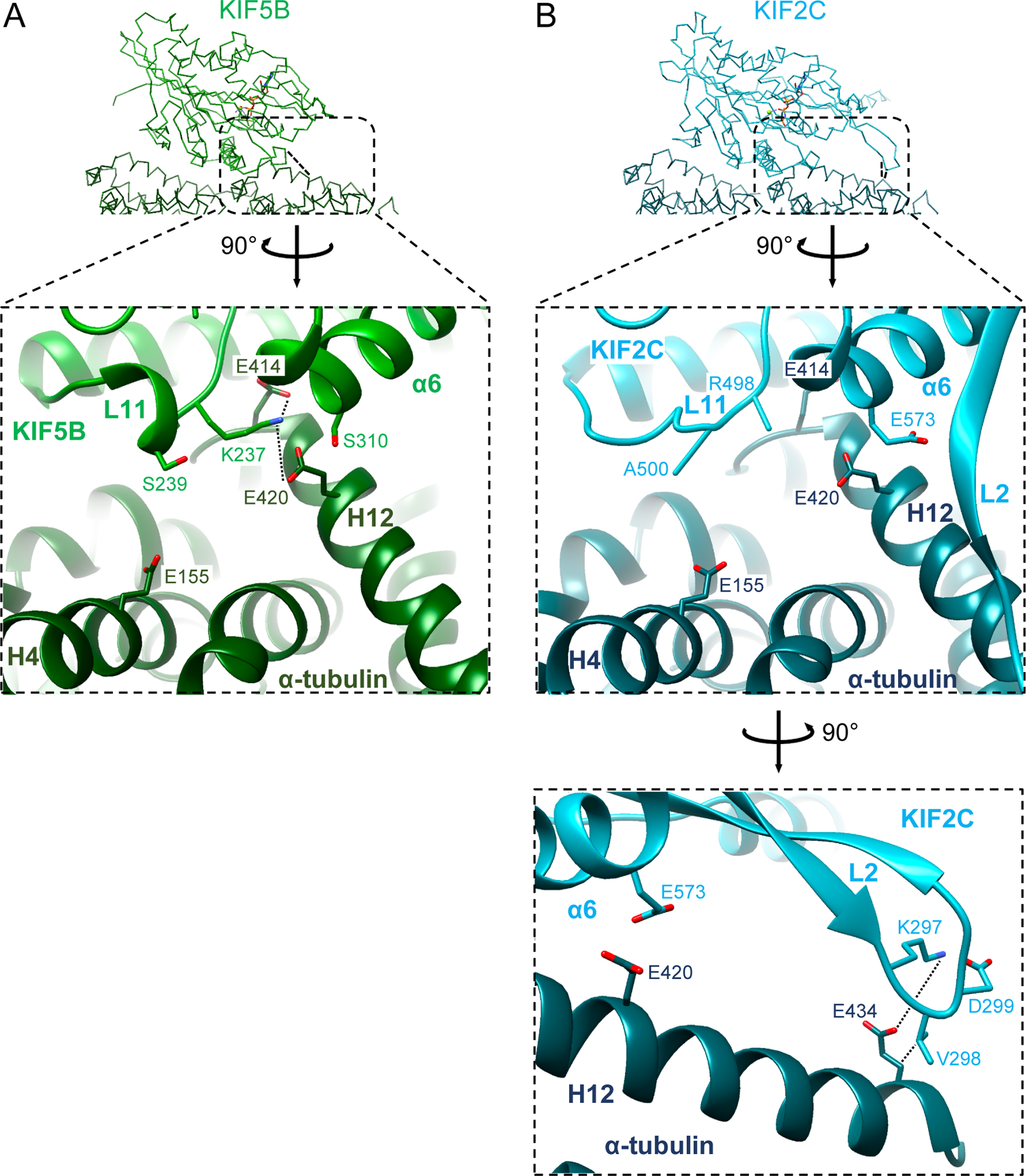
Microtubule binding interface of KIF5B and KIF2C at α-tubulin. **A.** KIF5B Lys237 in L11 interacts with α-tubulin Glu414 and Glu420 in H12. **B.** KIF2C Lys297 and Val298 in loop L2 KVD motif interact with α-tubulin Glu434 in H12. Arg498 was modeled as alanine in the reported structure (PDB ID: 5MIO). KVD motif residues Lys297, Val298, and Asp299 are displayed.

The former Arg267-mediated interaction is conserved in KIF5B through the corresponding residue Lys237 in L11 (Figure 4B; Figure 4–figure supplement 1A), but not conserved in KIF2C. KIF2C has Ala500 and Glu573 instead of Lys269 and Val343 of KLP-12, resulting in loss of the interaction (Figure 4B; Figure 4–figure supplement 1B). Instead, KIF2C takes another binding strategy through the KVD finger in loop L2 of KIF2C, as previously reported (Figure 4–figure supplement 1B) (Ogawa et al., 2004; Trofimova et al., 2018; Wang et al., 2017). The other interface between Lys269 of KLP-12 and Glu155 of α-tubulin is not conserved both in KIF5B and KIF2C. The counterpart residue is Ser239 in KIF5B, which is necessary for the kinesin-1-induced conformational change of microtubules to the growing GTP form (Shima et al., 2018) (Figure 4B; Figure 4–figure supplement 1A).

In summary, KLP-12 takes unique triangle contacts with α-tubulin conserved among kinesin-4. The different interfacial organizations of KLP-12 and KIF5B to α-tubulin result in the rotation of α-tubulin up to 4 degrees around Glu420 of α-tubulin (Figure 4C). On the other hand, although the interfaces between KLP-12 and KIF2C to α-tubulin are entirely different, the relative binding angle of KLP-12 and KIF2C with α-tubulin is consequently similar (Figure 4D). KLP-12, thus, induces a larger rotation of α-tubulin than KIF5B, which builds a composition of KIF2C similar to that of α-tubulin.

### Microtubule binding interface of kinesin-4 KLP-12 at β-tubulin

We next investigated the microtubule-binding interface of KLP-12 at β-tubulin by comparing it to the KIF5B and KIF2C structures superimposed using the kinesin motor domain (Figure 5; Figure 5–figure supplement 1). Helix α5 of KLP-12 and helix H12 of β-tubulin serve as the interface between KLP-12 and microtubules. Two arginine residues, Arg311 and Arg317, in α5 of KLP-12 form ionic interactions with Glu410 in H12 of β-tubulin (Figure 5A). These arginine residues are conserved in KIF5B (Figure 5B; Figure 1–figure supplement 2); however, they form intramolecular contacts with Glu157 in loop L8 of KIF5B instead of interacting with Glu420 (corresponding to Glu410 in porcine) of β-tubulin (Figure 5–figure supplement 1A), suggesting that the organization of α5 of KLP-12 or KIF5B gives rise to the difference.

**Figure 5.**
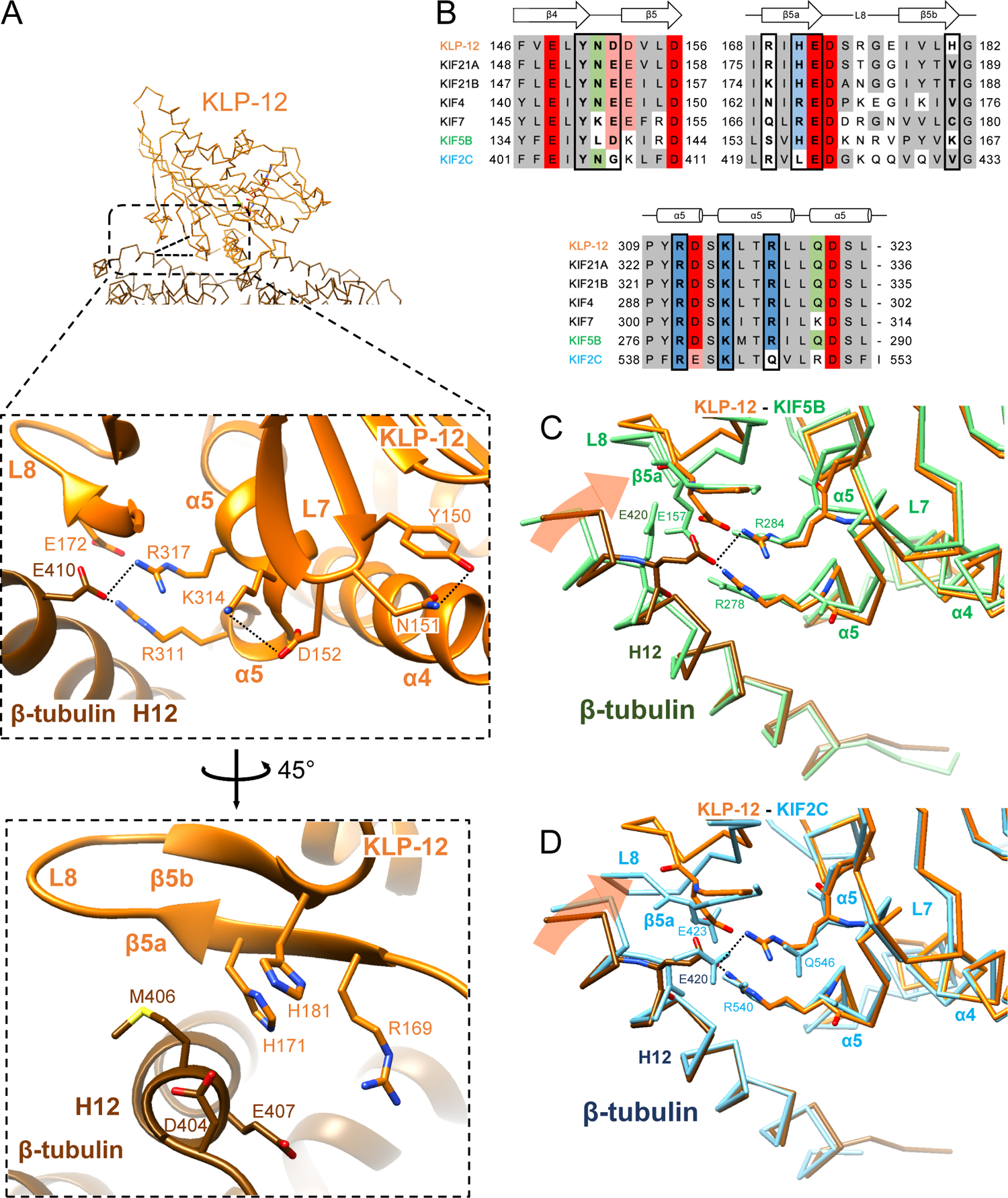
Microtubule binding interface of kinesin-4 KLP-12 at β-tubulin helix H12. (A) Close-up display of KLP-12 and β-tubulin interaction around β-tubulin helix H12 from the same view as upper panel (middle panel) and 45° rotated view (bottom panel). Glu410 of β-tubulin H12 interacts with Arg311 and Arg317 of KLP-12 helix α5. Tyr150 and Asn151 of KLP-12 form intramolecular interactions. The interfaces of KIF5B and KIF2C at β-tubulin are available in Figure 5 - figure supplement 1. (B) Sequence alignment with the secondary structure of the kinesin-4, KIF5B, and KIF2C residues at the interacting area. (C) Superimposition of Cα chain trace models of the KLP-12 complex (orange) and KIF5B complex (green) at kinesin around β-tubulin H12. (D) Superimposition of Cα chain trace models of the KLP-12 complex (orange) and KIF2C complex (cyan) at kinesin around β-tubulin H12.

**Figure 5 - figure supplement 1.**
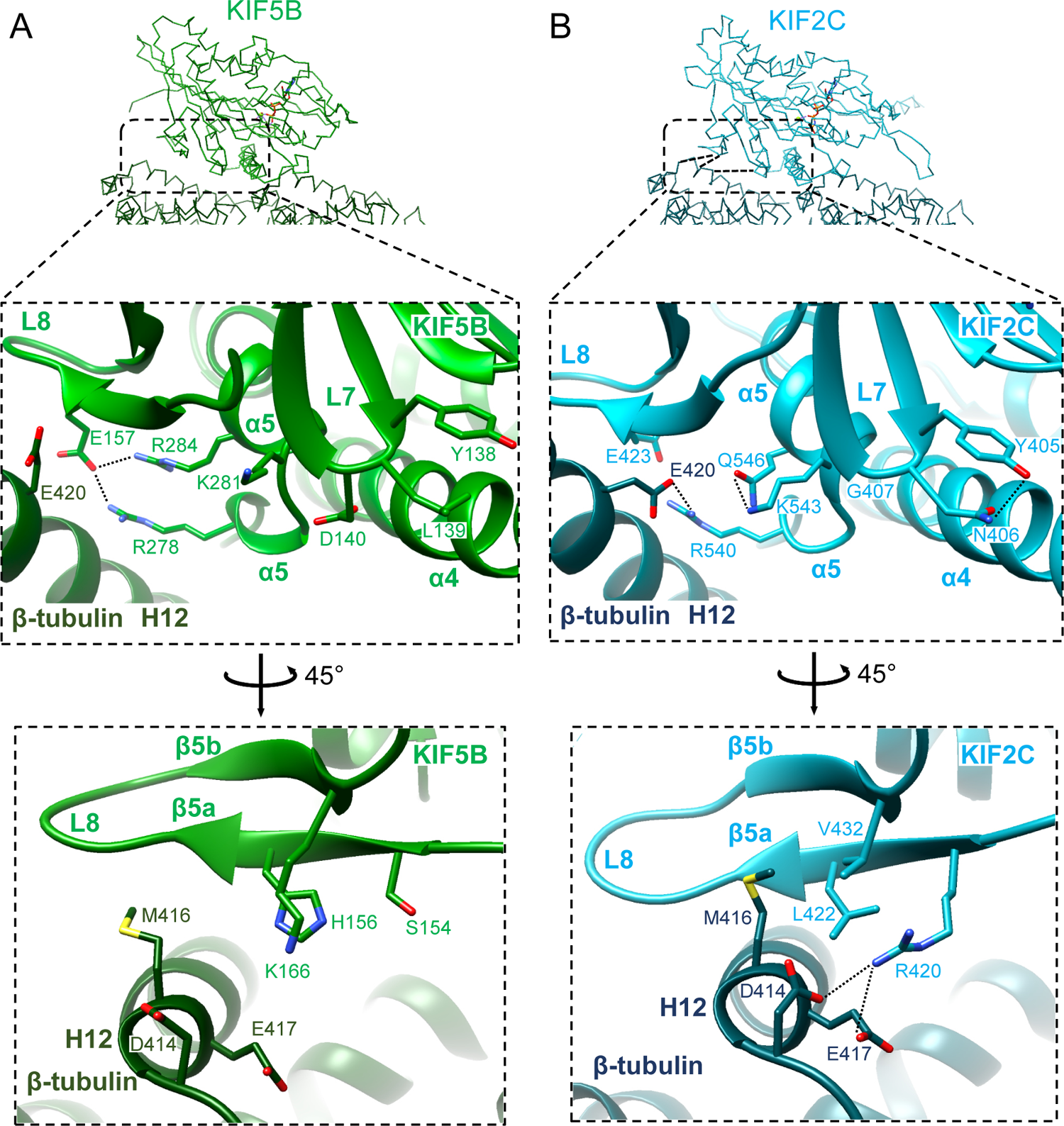
Microtubule binding interface of KIF5B and KIF2C at β-tubulin helix H12. **A.** KIF5B and β-tubulin interface. Glu420 (bovine Glu420 corresponds to Glu410 of porcine) of β-tubulin H12 has no interaction with KIF5B; instead, KIF5B Glu157 forms an intramolecular interaction with Arg278 and Arg284 of KIF5B α5. **B.** KIF2C and β-tubulin interaction. Glu420 (bovine Glu420 corresponds to Glu410 of porcine) of β-tubulin H12 interacts with Arg540 of KIF2C α5. Tyr405 and Asn406 of KIF2C form intramolecular interactions. In addition, KIF2C Arg420, which is supported by Leu422, interacts with Asp414 and Glu417 of β-tubulin H12.

To further investigate the interactions around α5 of KLP-12 and KIF5B, we carefully observed and found the intramolecular contact between Tyr150 and Asn151 and the resulting additional contact between Asp152 and Lys314 of KLP-12; Asn151 is highly conserved in kinesin-4 motors and KIF2C, but not in KIF5B (Figure 5B; Figure 1–figure supplement 2). The corresponding residue, Leu139 in KIF5B, did not form interaction, therefore Asp140 and Lys281 did not contact (Figure 5B; Figure 5–figure supplement 1A). The interaction within KLP-12 around Asn151 slightly rearranges the composition of α5 to generate intermolecular interaction between Glu410 of β-tubulin instead of intramolecular interaction with Glu172 of KLP-12. This interacting strategy is basically conserved in KIF2C (Figure 5B; Figure 5–figure supplement 1B).

In addition to this interaction, KIF2C has another intermolecular contact, which KLP-12 and KIF5B do not have. Arg420 in β5a of KIF2C forms an ionic interaction with Asp414 and Glu417 at the N-terminal side of H12 of β-tubulin (Figure 5–figure supplement 1B). This conformation of Arg420 is supported by Leu422 through hydrophobic contact with the side chain stem of Arg420. KLP-12 has His171 instead of Leu422 of KIF2C, showing the repulsive force keeping Arg169, which corresponds to residue Arg420 of KIF2C, away from the acidic residues on H12 (Figure 5A and B). KIF5B has Ser154 in the Arg420 position of KIF2C, resulting in no interaction with H12 of β-tubulin (Figure 5B; Figure 5–figure supplement 1A). These different types of interactions through H12 result in different rotation angles of β-tubulin. Compared to KIF5B, KLP-12 induces a ≃ 4 degree rotation of β-tubulin in the kinesin direction (Figure 5C). Comparison of KLP-12 and KIF2C generated further rotation of β-tubulin, resulting in the highest curvature of tubulin to destabilize the microtubule end (Figure 5D).

### Distinct curvature of tubulin-dimer induced by different kinesin motor domains

As described above, the different types of kinesin motor domains arrange the different kinesin–tubulin interfaces to produce the distinct curvature of tubulin-dimer. To elucidate how kinesin-4 family KLP-12 affects the overall conformation of the tubulin dimer, kinesin–tubulin complex structures were superimposed on α-tubulin and compared (Figure 6A-C). As a result, obvious differences in β-tubulin positions or rotations were observed among the three kinesins. The tubulin dimer with KIF5B, which generates a glowing plus-end, forms the straightest conformation. The tubulin dimer with KIF2C, which is a plus-end destabilizing kinesin, forms the most curved conformation, 6 degrees larger than that of KIF5B. Intriguingly, KLP-12 induces intermediate curvature of the tubulin dimer, with the α- and β-tubulin angles being 3 degrees more curved than KIF5B.

**Figure 6.**
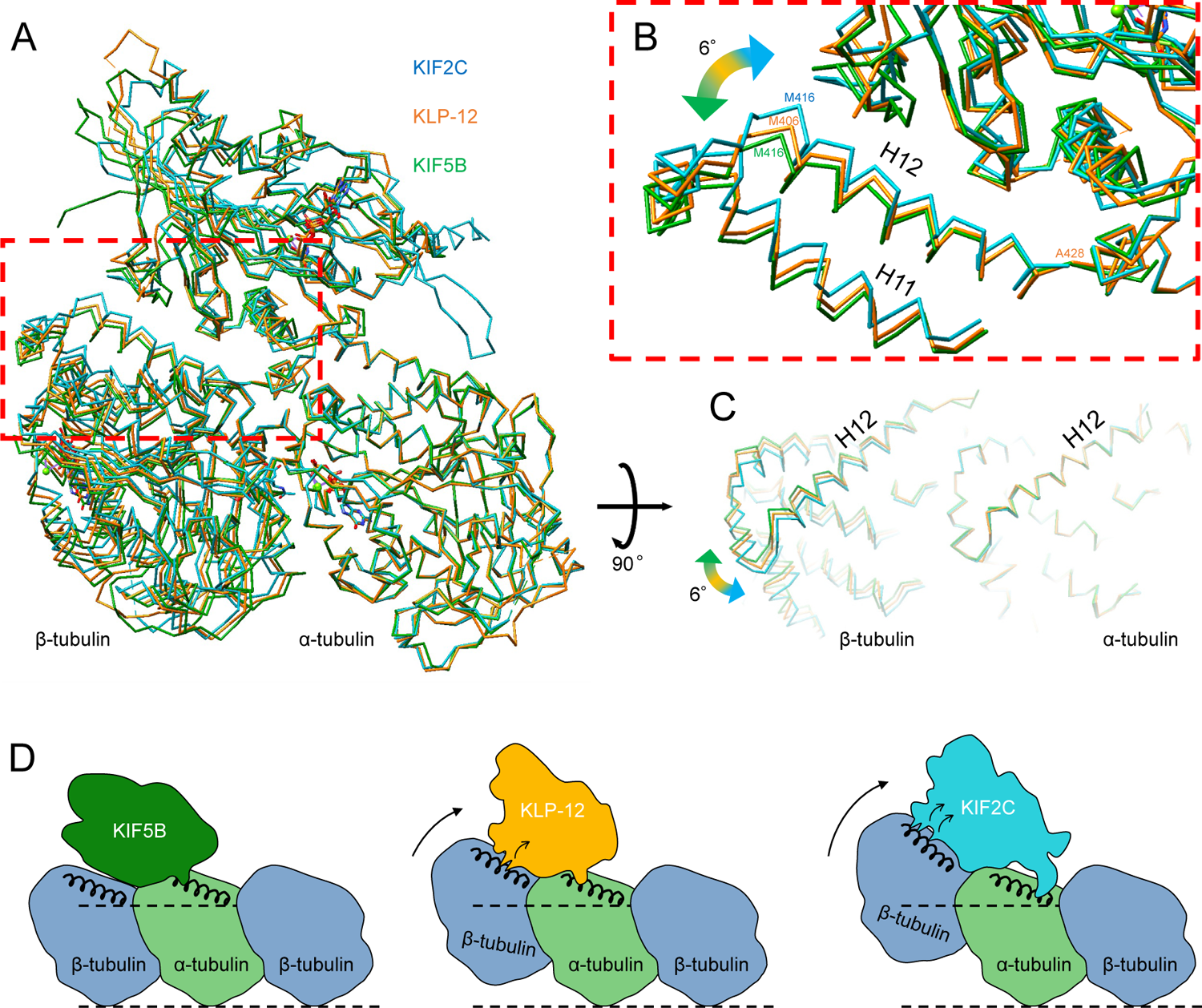
Microtubule plus-end control mechanism by kinesin-4. (A) Superimposition of KLP-12, KIF5B, and KIF2C complexes at α-tubulin. (B) Close-up view of superimposed models at the β-tubulin interaction surface. (C) Tubulin dimer interface view from kinesin side. (D) Models of the microtubule plus-end curving mechanism by kinesins. The kinesin-1 family protein KIF5B does not affect the curvature of microtubules to maintain the conformation of GTPs. Kinesin-4 family protein KLP-12 slightly curves the plus-end by interacting with the mid-portion of β-tubulin H12 to stabilize the plus-end. Kinesin-13 family protein KIF2C strongly curves plus-end by interacting with the N-terminal portion of β-tubulin H12 to destabilize the microtubule.

## DISCUSSION

This study investigated the molecular mechanism of microtubule dynamics inhibition by kinesin-4 motor *C. elegans* KLP-12 using genetic, biophysical, biochemical, and structural analyses. *C. elegans* genetics clearly illustrated the role of KLP-12 in regulating the length of axons by inhibiting microtubule dynamics. Similar to KIF21A/B, the motor domain and the tail domain are required to precisely control neurite length (van der Vaart et al., 2013; van Riel et al., 2017). Biophysical analyses elucidated the plus-end directed motility of KLP-12 and the inhibitory effect of microtubule dynamics by KLP-12. Biochemical analyses also showed that KLP-12 is similarly active on both the microtubule lattice and plus-end. Similar binding abilities and ATPase activities of KLP-12 on microtubules and tubulin dimers were detected. Structural studies demonstrated kinesin-4-specific residues at the microtubule-binding interface of KLP-12, adequately increasing the curvature of tubulin-dimers. It might enable the precise curvature of microtubule plus-ends to inhibit microtubule dynamics.

To evaluate the proper regulation of microtubule dynamics, we utilized axon length in *C. elegans* neurons. Defect of the motor domain or the tail domain resulted in the excessive elongation of axons, whereas the overexpression of KLP-12 strikingly shortened axons, concluding the role of KLP-12 in inhibiting microtubule dynamics (Figure 1D-G). We should note here the phenotypic difference between the motor and tail defects; the motor defect made the axon wavy and thinly, whereas the tail defect did not change the thickness of axons (Figure 1D). There are two possibilities: (I) motor function will be required for proper axon development, or (II) the tail domain without the motor will disable proper axon development. Future studies are required to elucidate this mechanism.

Steady state ATPase assays of the KLP-12 motor domain in the presence of microtubules or tubulin dimers elucidated the ATPase activities with comparable affinities to microtubules and tubulin dimers. Although kinesin-13 expresses stronger binding of tubulin dimers over microtubules, other kinesin motors bind strongly to microtubules over tubulin dimers, including Xklp1 or KIF19A (Bringmann et al., 2004; Hunter et al., 2003; Wang et al., 2016). Therefore, preferential binding to the plus-end of microtubules is one of the important features of KLP-12 to turn the pealed end into a workspace. However, since ATP hydrolysis detaches KLP-12 from the microtubule ends, the tail domain should be required to tether itself to microtubules to continuously stabilize the plus ends.

From the crystal structural analysis, we identified the specific interactions of KLP-12 for α- and β-tubulins (Figure 4; Figure 5). The contributing residues are mostly conserved among kinesin-4, suggesting the conserved structural mechanisms. KLP-12 interacts with α-tubulin through the triangle contacts at the N-terminal side of H12 and H4, and β-tubulin at the mid-portion of H12. Kinesin-1 makes single contact with α-tubulin at the N-terminal side of α-tubulin H12 but no contact with β-tubulin. Kinesin-13 binds to α-tubulin at the C-terminal side of H12 through the KVD finger and to β-tubulin at the N-terminal side of H12, in addition to the mid-portion contact. These interactions are summarized in the schematic illustrations in Figure 6D.

Kinesin-1 cannot bend the protofilament and allows microtubules to polymerize, whereas kinesin-13 strongly bends the protofilament to depolymerize microtubules (Ogawa et al., 2004; Shima et al., 2018). KLP-12 expresses the mild effect of protofilament bending, enabling the inhibition of both the polymerization and depolymerization of the microtubules. As summarized previously, changes in the curvature of the peeled plus-ends of microtubules are fundamental to microtubule dynamics (Brouhard and Rice, 2014); less curved plus-end polymerizes the microtubule, more curved plus-end shortens the microtubule, and middle curved plus-end stabilizes the microtubule. In addition to the reported structures of KIF5B with tubulin and KIF2C with tubulin representing polymerizing and depolymerizing plus-end of a microtubule, KLP-12 complexed with tubulin structure filled the missing piece of the structural information of stabilizing plus-end of a microtubule. The structural model might explain the molecular mechanism of distinct effects on the microtubule dynamics among kinesin-4 motors by controlling the bending angle; slightly more curving inhibits plus-end directed motility along the microtubule lattice, inhibits the polymerization of microtubules, and accelerates the destabilization of microtubules, possibly represented by KIF7. The other domains, including the tail, would further intricately arrange the bending effects of motor domains.

In summary, our study provides important structural clues regarding a novel molecular mechanism by which kinesin-4 inhibits microtubule dynamics. This structural model is consistent with the previously reported effects of microtubule elongation or destabilization by the motor domains of various kinesin superfamily proteins.

## Materials and methods

### Key resources table

### List of strains used in this study

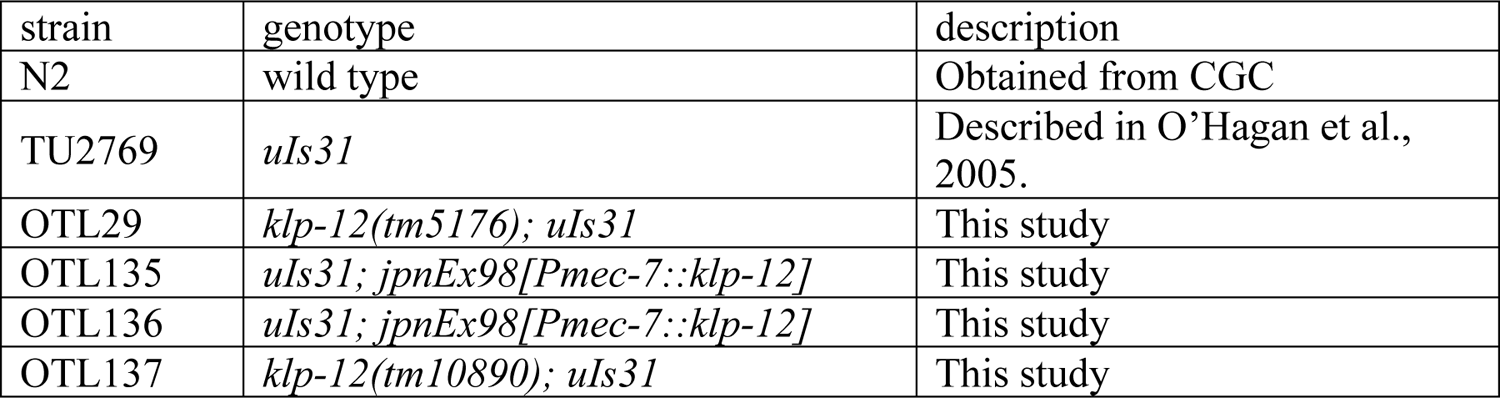

### List of protein expression constructs used in this study

| ID | construct | vector | tag (N-terminal) | tag (C-terminal) |
| --- | --- | --- | --- | --- |
| RN349 | KLP-12-LZ-GFP | pET21B | none | 6x His |
| RN107 | KLP-12(M) | pGEX-6P | GST | none |
| RN84 | DARPin | pETDUET-1 | none | 6x His |
| RN148 | KLP-12-DARPin | pGEX-6P | GST | none |

### The amino acid sequence of constructs used in this study

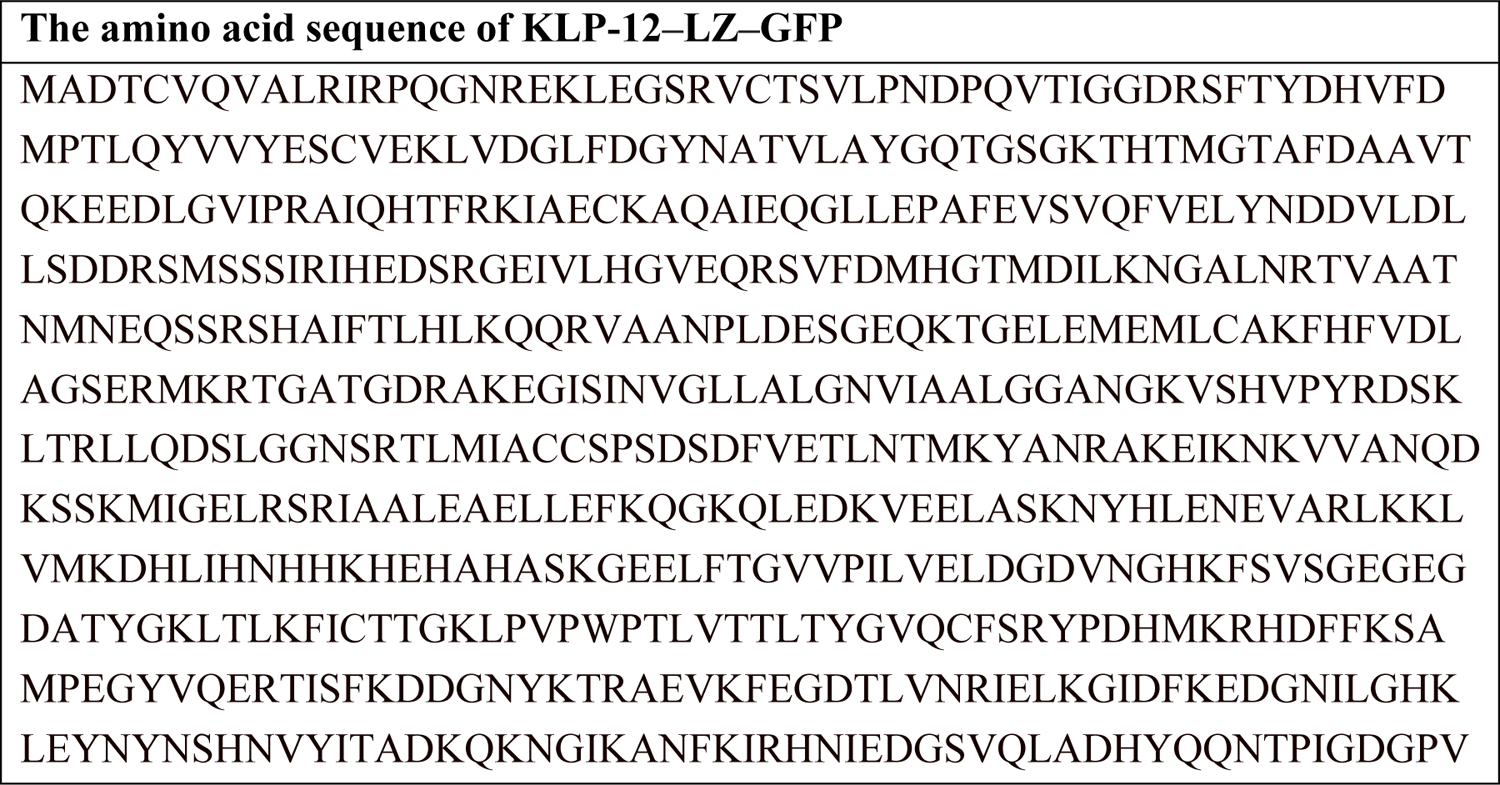
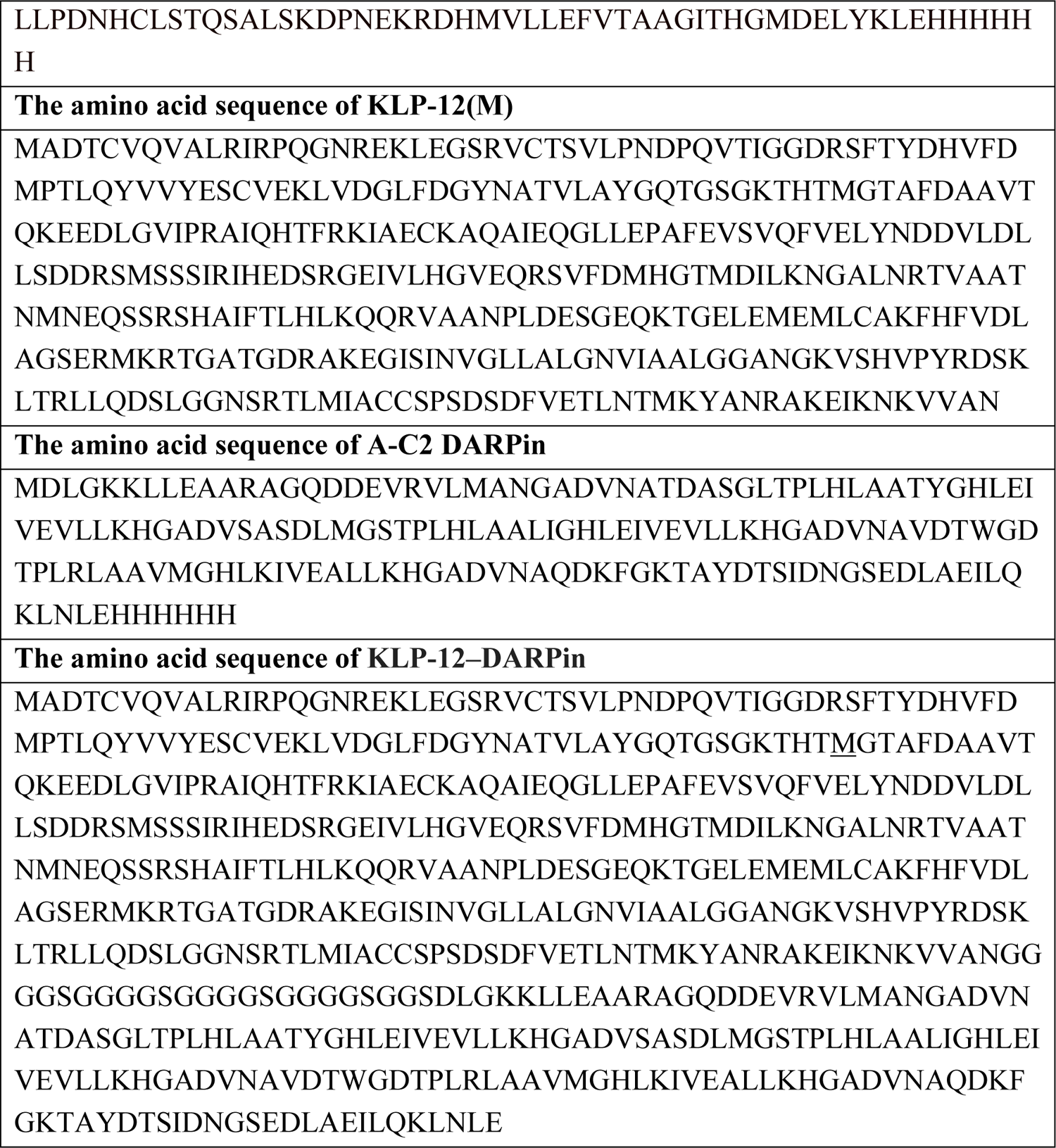

#### C. *elegans* experiments

The strains used in this study are described in Key resources table. *C. elegans* strains were maintained as described previously (Brenner, 1974). Some strains and OP50 feeder bacteria were obtained from the *C. elegans* genetic center (CGC) (Minneapolis, MN, USA). *klp-12(tm5176)* and *klp-12(tm10890)* were obtained from the National BioResource Project (NBRP, Tokyo, Japan). Transformation of *C. elegans* was performed by DNA injection as described (Mello et al., 1991). *uIs31[Pmec-17::GFP]* was used to visualize the morphology of mechanosensory neurons (O’Hagan et al., 2005). Worms were fixed using 0.25 mM levamisol (Sigma), 5% agarose pads and 0.1 μm polystylene beads (Polysciences) as described (Niwa, 2017). Worms were observed by an LSM800 confocal microscope system equipped with a x20 Plan-Apochromat objective lens (NA0.8) (Carl Zeiss).

#### Cloning of *klp-12* cDNA

Total worm cDNA was a kind gift from Dr. Kota Mizumoto (University of British Columbia). First, we obtained the full-length *klp-12* cDNA fragment by PCR performed using KOD FX neo (TOYOBO, Tokyo, JAPAN). The first PCR was performed with 5’-gtgaaaATGGCGGACACTTGTGTGC-3’ and 5’-ATTACAGAAAAGTGAAAAGGGGTACAAGTG-3’ primers, and then the second PCR was performed with 5’-gtgaaaATGGCGGACACTTGTGTGC-3’ and 5’-GACAGCATTTGATTTCCAGAATCCGAC-3’. We fully sequenced the fragment and confirmed the sequence of full-length *klp-12* cDNA (NM_001028178). However, we found that the 3’ latter half of the *klp-12* cDNA had toxicity in bacteria and could not be inserted into vectors such as pBluescript, pEGFPN1 and pFastbac. Toxicity-resistant strains such as NEB 5-alpha F’I^q^ (NEB) and ABLE K (Agilent) could not compromise toxicity. Then, based on the sequence data, GeneArt (Thermo Fisher Scientific) was used to synthesize full-length *klp-12* cDNA, which was codon-optimized for *C. elegans*.

### Tubulin preparation

Tubulin was purified from porcine brains following a previously reported method (Castoldi and Popov, 2003).

### Plasmid construction

For preparation of the KLP-12(M) construct, the coding region of KLP-12 (1–365) was further amplified by PCR with an overlapping sequence with pGEX-6P and cloned immediately after the PreScission site of the pGEX-6P vector by Hi-Fi DNA assembly (NEB).

For preparation of the KLP-12–LZ–GFP construct, the N-terminal end of GFP connected to the C-terminal end of KLP-12(1-393) with the leucine zipper (LZ) was amplified by PCR by adding a LZ sequence (Yue et al., 2018) and a C-terminal 6x His tag and cloned into the pET21B vector by Hi-Fi DNA assembly (NEB).

For preparation of the DARPin construct, the coding sequence of A-C2 DARPin was synthesized by the manufacturer (IDT gBlocks). The DNA fragment was amplified by PCR by inserting a C-terminal 6x His tag and cloned into the pETDUET-1 vector by Hi-Fi DNA assembly (NEB).

For preparation of the KLP-12–DARPin construct, the N-terminal end of DARPin connected to the C-terminal end of KLP-12(M) with the (G_4_S)_4_-GGS linker was amplified by PCR by adding a linker sequence and inserted into the pGEX-6P vector by Hi-Fi DNA assembly (NEB).

All constructs used in the study are listed in Key resources table.

### Protein expression and purification

To obtain the KLP-12(M) constructs, the *Escherichia coli* strain BL21(DE3) harboring the plasmid encoding the KLP-12 (1–365) fragment was cultured in LB medium containing 50 μg/ml ampicillin at 37 °C until OD600 > 0.4. Protein expression was induced by 0.2 mM IPTG at 18 °C overnight. The cells were harvested by centrifugation and resuspended in 20 mM Tris-HCl pH 8.0 and 171 mM NaCl. The cell suspension was harvested by centrifugation and stored at −80 °C. The cells were resuspended in 50 mM HEPES-KOH pH 7.5, 400 mM KCl, 10% glycerol, 2 mM MgCl_2_, 5 mM β-mercaptoethanol, and protease inhibitor (0.7 μM leupeptin, 2 μM pepstatin A, 1 mM PMSF, and 2 mM benzamidine). Lysed cells were disrupted using sonication, and the cell debris was removed by centrifugation using an Avanti JXN-30 centrifuge (Beckman Coulter) with a JA-30.50Ti rotor at 80,000 × g. The supernatant was loaded onto 2 ml Glutathione Sepharose^TM^ 4B (GE Health care) for affinity chromatography, followed by 3C protease cleavage of glutathione S-transferase (GST). After concentration using an Amicon Ultra concentrator (Merck Millipore; 10 kDa MWCO), the target protein was further purified on a Superdex 200 Increase 10/300 GL size exclusion column (GE Health care) in 20 mM HEPES-KOH pH 7.5, 150 mM NaCl, 2.5 mM MgCl_2_, and 1 mM DTT. Peak fractions were pooled, concentrated with an Amicon Ultra 10 kDa MWCO concentrator until 10 mg/ml, flash frozen by liquid nitrogen, and stored at −80 °C until use.

To obtain KLP-12–LZ–GFP, the same protocol as for KLP-12(M) was used until the cell suspension was harvested and stored at −80 °C. The cells were resuspended in wash buffer (50 mM HEPES-KOH pH 7.5, 400 mM KCl, 10% glycerol, 2 mM MgCl_2_, 5 mM imidazole, 5 mM β-mercaptoethanol, and protease inhibitor) and disrupted using sonication. The cell debris was removed by centrifugation using an Avanti JXN-30 centrifuge (Beckman Coulter) with a JA-30.50Ti rotor at 80,000 × g. The supernatant was loaded on a 2 ml HIS-Select^TM^ Nickel Affinity Gel (Merck) and washed with wash buffer. Bound protein was eluted with 50 mM HEPES-KOH pH 7.5, 400 mM KCl, 10% glycerol, 2 mM MgCl_2_, 300 mM imidazole, 5 mM β-mercaptoethanol, and protease inhibitor. After concentration using an Amicon Ultra 10 kDa MWCO concentrator, the target protein was further purified on a Superdex 200 Increase 10/300 GL size exclusion column (GE Health care) in 20 mM HEPES-KOH pH 7.5, 150 mM NaCl, 2.5 mM MgCl_2_, and 1 mM DTT. Peak fractions were pooled, and the target protein was further purified on a anion-exchange Mono Q column (GE Health care) with a linear gradient of NaCl in A buffer (20 mM HEPES-KOH pH 7.5, 50 mM NaCl, 2.5 mM MgCl_2_, and 1 mM DTT) and B buffer (20 mM HEPES-KOH pH 7.5, 1 M NaCl, 2.5 mM MgCl_2_, and 1 mM DTT). Peak fractions were pooled, concentrated and buffer changed with an Amicon Ultra 10 kDa MWCO concentrator into A buffer, flash frozen by liquid nitrogen and stored at −80 °C until use.

To obtain the DARPin constructs, the *E. coli* strain BL21(DE3) harboring the plasmid encoding the A-C2 DARPin fragment was cultured in LB medium containing 100 μg/ml ampicillin at 37 °C until OD600 > 0.4. Protein expression was induced by 0.1 mM IPTG at 30 °C overnight. The cells were harvested by centrifugation and resuspended in 20 mM Tris-HCl pH 8.0 and 171 mM NaCl. The cell suspension was harvested by centrifugation and stored at −80 °C. The cells were resuspended in wash buffer (50 mM Tris-HCl pH 8.0, 300 mM NaCl, 5 mM imidazole, 5 mM β-mercaptoethanol, and protease inhibitor) and disrupted using sonication. The cell debris was removed by centrifugation using an Avanti JXN-30 centrifuge (Beckman Coulter) with a JA-30.50Ti rotor at 80,000 × g. The supernatant was loaded on a 2 ml HIS-Select^TM^ Nickel Affinity Gel (Merck) and washed with wash buffer. Then, a high salt wash (50 mM Tris-HCl pH 8.0, 1 M NaCl, 5 mM imidazole, 5 mM β-mercaptoethanol, and protease inhibitor) was performed followed by a low salt wash (50 mM Tris-HCl pH 8.0, 10 mM KCl, 5 mM imidazole, 5 mM β-mercaptoethanol, and protease inhibitor). Bound protein was eluted with 50 mM Tris-HCl pH 8.0, 100 mM KCl, 250 mM imidazole, 5 mM β-mercaptoethanol, and protease inhibitor. After concentration using an Amicon Ultra 10 kDa MWCO concentrator, the target protein was further purified on a Superdex 75 10/300 GL size exclusion column (GE Health care) in 20 mM HEPES-KOH pH 7.5, 150 mM NaCl, 2.5 mM MgCl_2_, and 1 mM DTT. Peak fractions were pooled, concentrated with an Amicon Ultra 10 kDa MWCO concentrator, flash frozen by liquid nitrogen and stored at −80 °C until use.

To obtain KLP-12–DARPin, the same protocol as for KLP-12(M) was used for protein expression and purification, except the use of 100 μg/ml ampicillin for LB medium and 0.1 mM IPTG for protein expression induction.

### Observation of MT dynamics

The in vitro MT observation was performed as reported before (Al-Bassam, 2014). Microscopy slides and precleaned glass coverslips were used to assemble a flow chamber using double-sided tape. The chamber was treated with 0.5 mg/ml PLL-PEG-biotin (Surface Solutions, Switzerland) in BRB80 buffer (80 mM K-PIPES, pH 6.8, 1 mM MgCl2, and 1 mM EGTA) for 5 min. After washing the chamber with BRB80 buffer, it was incubated with 0.5 mg/ml Streptavidin for 5 min. Short MT seeds were prepared using 1.5 μM tubulin mix containing 50% biotin-tubulin and 50% AZdye647-tubulin with 1 mM GMPCPP at 37 °C for 30 min. The polymerized MTs were pelleted by ultracentrifuge for 5 min. The pellet was resuspended by BRB80 supplemented with 10% glycerol and fragmented by pipetting. Resultant seeds were aliquoted and snapfrozen by liquid N2. The seeds were melted on 37 °C heat block immediately before the use and diluted by BRB80. Seeds were attached on the coverslips using biotin-avidin links and incubated with assay buffer (80 mM K-PIPES, pH 6.8, 1 mM MgCl2, and 1 mM EGTA, 0.5% Pluronic F127, 0.1 mg/ml BSA, 0.1 mg/ml biotin-BSA, 0.2 mg/ml k-casein).

The in vitro reaction mixture consisting of 10 μM tubulin, in assay buffer supplemented with 1 mM GTP, oxygen scavenging system composed of Trolox/PCD/PCA, 2 mM ATP, and the specified amount of KLP-12–LZ–GFP was added to the flow chamber. During the experiments the samples were maintained at 30 °C. An ECLIPSE Ti2-E microscope equipped with a CFI Apochromat TIRF 100XC Oil objective lens, an Andor iXion life 897 camera and a Ti2-LAPP illumination system (Nikon, Tokyo, Japan) was used to observe single molecule motility. NIS-Elements AR software ver. 5.2 (Nikon) was used to control the system.

### Single molecule observation

TIRF assay was performed as described before (Chiba et al., 2019).

To polymerize Taxol-stabilized microtubules labeled with biotin and AZDye647, 30 μM unlabeled tubulin, 1.5 μM biotin-labeled tubulin and 1.5 μM AZDye647-labeled tubulin were mixed in BRB80 buffer supplemented with 1 mM GTP and incubated for 15 min at 37 °C. Then, an equal amount of BRB80 supplemented with 40 μM taxol was added and further incubated for more than 15 min. The solution was loaded on BRB80 supplemented with 300 mM sucrose and 20 μM taxol and ultracentrifuged at 100,000 g for 5 min at 30 °C. The pellet was resuspended in BRB80 supplemented with 20 μM taxol. Polymerized microtubules were flowed into streptavidin adsorbed flow chambers and allowed to adhere for 5–10 min. Unbound microtubules were washed away using assay buffer supplemented with taxol. Purified motor protein was diluted to indicated concentrations in the assay buffer suppelemented with 2 mM ATP and oxygen scavenging system composed of Trolox/PCD/PCA. Then, the solution was flowed into the glass chamber. The sample was observed by the TIRF system as described above.

### ATPase activity

ATPase activity was measured using the EnzChek^TM^ Phosphate Assay Kit (Molecular Probes) and DS-11+ spectrophotometer (DeNovix). 45.6 μM tubulin was polymerized in PEM buffer (0.1 M PIPES-KOH pH 6.8, 1 mM EGTA, 1 mM MgCl_2_) supplemented with 1 mM GTP and 4% DMSO (PEM-GTP buffer) at 37 °C for 30 min. Each concentration used in the experiment was diluted with PEM-GTP buffer. When measuring tubulin, the polymerization step was omitted. The reactions were performed in 5 mM HEPES-KOH pH 7.5, 5 mM potassium acetate, 200 μM MESG, 1.5 U/ml PNP, 10 μM Taxol, and all assays were performed at 30 °C. When measuring as tubulin, Taxol was not added.

ATPase activity: After 1 mM ATP was added and the absorbance became stable, 1 μM KLP-12(M) was added, and the absorbance was measured every 5 seconds. It was calculated by subtracting the absorbance of the control measured without KLP-12(M). The experiment was performed 3 times. Values for V_max_ and K_m_ were obtained by Lineweaver-Burk plots of ATPase activity versus microtubule or tubulin concentration using Microsoft Excel (R^2^=0.996 or 0.990, respectively).

Basal ATPase activity: PEM-GTP buffer was used instead of microtubules (tubulin), no Taxol was added, and measurements were taken at 0.06 U/ml PNP to delay the reaction. After 1 mM ATP was added and the absorbance became stable, 1 μM KLP-12(M) was added, and the absorbance was measured every 5 seconds. It was calculated by subtracting the absorbance of the control measured without KLP-12(M). The experiment was performed 3 times.

### Size exclusion chromatography analyses

KLP-12–DARPin with tubulin or KLP-12(M), DARPin, and tubulin were analyzed on a Superdex 200 Increase 10/300 GL column equilibrated with 20 mM PIPES-KOH pH 6.8, 50 mM KCl, 1 mM MgCl_2_, 0.5 mM EGTA, and 10 μM AMP-PNP. Following gel filtration, the proteins were precipitated with acetone, separated by SDS–PAGE, and visualized by Coomassie staining.

### Crystallization and structure determination

Then, 0.6 mg/ml tubulin–KLP-12–DARPin complex, which was purified by size exclusion chromatography, was crystallized at 20 °C by vapor diffusion in crystallization buffer containing 0.2 M ammonium acetate, 0.1 M HEPES pH 7.2–7.8, 22–26% polyethylene glycol (PEG) 3350. Crystals were harvested by Litholoop and transferred into cryoprotectant solution containing 0.2 M ammonium acetate, 0.1 M HEPES pH 7.4, 25% PEG 3350, and 20% glycerol and then flash-frozen in liquid nitrogen. All diffraction datasets were collected at the BL32XU beamline in a synchrotron facility SPring-8 equipped EIGER 9M detector (DECTRIS Ltd) at −180 °C with wavelengths of 1.00000 Å followed by a ZOO system (Hirata et al., 2019). Loop-harvested microcrystals were identified using raster scan and analysis by SHIKA (Hirata et al., 2019). Small wedge data, each consisting of 10°, were collected from single crystals, and the collected datasets were processed automatically using KAMO (Yamashita et al., 2018) with XDS (Kabsch, 2010a), followed by hierarchical clustering analysis using the correlation coefficients of the normalized structure amplitudes between datasets. Finally, a group of outlier-rejected datasets were scaled and merged using XSCALE (Kabsch, 2010b). The structure was determined by molecular replacement with the program PHASER (McCoy et al., 2007) using the crystal structure of tubulin dimer (PDB ID: 5MIO) as ensamble 1 and the homology model of KLP-12 generated by SWISS-MODEL (Waterhouse et al., 2018) with DARPin (PDB ID: 5MIO) as ensamble 2. The electron density map and the structural model were iteratively refined and rebuilt using PHENIX and COOT (Liebschner et al., 2017). The Ramachandran statistics are 92.5% and 7.5% in the favored and allowed regions of the Ramachandran plots, respectively, and 0% are outliers. Data collection and refinement statistics are summarized in Table 1. All molecular graphics were prepared by using UCSF Chimera (Pettersen et al., 2004).

## Acknowledgements

We thank K. Chin, T. Setsu, and Y. Sakihama for assistance and other colleagues for discussions. This work was supported by the Platform Project for Supporting Drug Discovery and Life Science Research (Basis for Supporting Innovative Drug Discovery and Life Science Research (BINDS)) from AMED under Grant Number JP21am0101070. We acknowledge support from the Japan Society for the Promotion of Science (KAKENHI; 19K07246 to T. I., and 19H03396 (R.N.)), AMED-CREST from the Japan Agency for Medical Research and Development (JP21gm0810013 to R. N.), JST (Moonshot R&D)(Grant Number JPMJMS2024), the Takeda Science Foundation to T. I. and R. N, the Mochida Memorial Foundation for Medical and Pharmaceutical Research to T. I. and R. N., the Uehara Memorial Foundation to R. N., Bristol-Myers Squibb to R. N., and the Hyogo Science and Technology Association to R. N.

## AUTHOR CONTRIBUTIONS

S. T., T. I., S. N. and R. N. conceived the project. S. T., T. I., Y. S. H., N. S., H. S., H. O., T. S., E. N., S. K., and S. M. performed DNA construction, protein purification, and biochemical analyses. J. N., T. K., and S. N. performed *C. elegans* and TIRF assays. S. T., J. N., T. I., S. N., and R. N. prepared the figures. All authors discussed the results, and S. T., T. I., S. N., and R. N. wrote the manuscript.

## Data availability

Data generated during this study are included in the manuscript and supplemental information. Protein coordinates and structural factors have been deposited in the RCSB Protein Data Bank (https://www.rcsb.org) with accession code PDB: ….

## REFERENCES

1. Ahmad S, Pecqueur L, Dreier B, Hamdane D, Aumont-Nicaise M, Plückthun A, Knossow M, Gigant B. 2016. Destabilizing an interacting motif strengthens the association of a designed ankyrin repeat protein with tubulin. Sci Rep 6:28922. doi:10.1038/srep28922

2. Al-Bassam J. 2014. Reconstituting dynamic microtubule polymerization regulation by TOG domain proteins. Methods Enzymol 540:131–148. doi:10.1016/B978-0-12-397924-7.00008-X

3. Barrett JC, Hansoul S, Nicolae DL, Cho JH, Duerr RH, Rioux JD, Brant SR, Silverberg MS, Taylor KD, Barmada MM, Bitton A, Dassopoulos T, Datta LW, Green T, Griffiths AM, Kistner EO, Murtha MT, Regueiro MD, Rotter JI, Schumm LP, Steinhart AH, Targan SR, Xavier RJ, NIDDK IBD Genetics Consortium, Libioulle C, Sandor C, Lathrop M, Belaiche J, Dewit O, Gut I, Heath S, Laukens D, Mni M, Rutgeerts P, Van Gossum A, Zelenika D, Franchimont D, Hugot J-P, de Vos M, Vermeire S, Louis E, Belgian-French IBD Consortium, Wellcome Trust Case Control Consortium, Cardon LR, Anderson CA, Drummond H, Nimmo E, Ahmad T, Prescott NJ, Onnie CM, Fisher SA, Marchini J, Ghori J, Bumpstead S, Gwilliam R, Tremelling M, Deloukas P, Mansfield J, Jewell D, Satsangi J, Mathew CG, Parkes M, Georges M, Daly MJ. 2008. Genome-wide association defines more than 30 distinct susceptibility loci for Crohn’s disease. Nat Genet 40:955–962. doi:10.1038/ng.175

4. Bianchi S, van Riel WE, Kraatz SHW, Olieric N, Frey D, Katrukha EA, Jaussi R, Missimer J, Grigoriev I, Olieric V, Benoit RM, Steinmetz MO, Akhmanova A, Kammerer RA. 2016. Structural basis for misregulation of kinesin KIF21A autoinhibition by CFEOM1 disease mutations. Sci Rep 6:30668. doi:10.1038/srep30668

5. Bieling P, Telley IA, Surrey T. 2010. A minimal midzone protein module controls formation and length of antiparallel microtubule overlaps. Cell 142:420–432. doi:10.1016/j.cell.2010.06.033

6. Brenner S. 1974. The genetics of Caenorhabditis elegans. Genetics 77:71–94.

7. Bringmann H, Skiniotis G, Spilker A, Kandels-Lewis S, Vernos I, Surrey T. 2004. A kinesin-like motor inhibits microtubule dynamic instability. Science 303:1519–1522. doi:10.1126/science.1094838

8. Brouhard GJ, Rice LM. 2014. The contribution of αβ-tubulin curvature to microtubule dynamics. J Cell Biol 207:323–334. doi:10.1083/jcb.201407095

9. Castoldi M, Popov AV. 2003. Purification of brain tubulin through two cycles of polymerization-depolymerization in a high-molarity buffer. Protein Expr Purif 32:83–88. doi:10.1016/S1046-5928(03)00218-3

10. Chang Q, Nitta R, Inoue S, Hirokawa N. 2013. Structural basis for the ATP-induced isomerization of kinesin. J Mol Biol 425:1869–1880. doi:10.1016/j.jmb.2013.03.004

11. Cheng L, Desai J, Miranda CJ, Duncan JS, Qiu W, Nugent AA, Kolpak AL, Wu CC, Drokhlyansky E, Delisle MM, Chan W-M, Wei Y, Propst F, Reck-Peterson SL, Fritzsch B, Engle EC. 2014. Human CFEOM1 mutations attenuate KIF21A autoinhibition and cause oculomotor axon stalling. Neuron 82:334–349. doi:10.1016/j.neuron.2014.02.038

12. Chiba K, Takahashi H, Chen M, Obinata H, Arai S, Hashimoto K, Oda T, McKenney RJ, Niwa S. 2019. Disease-associated mutations hyperactivate KIF1A motility and anterograde axonal transport of synaptic vesicle precursors. Proc Natl Acad Sci U S A 116:18429–18434. doi:10.1073/pnas.1905690116

13. Desai A, Verma S, Mitchison TJ, Walczak CE. 1999. Kin I kinesins are microtubule-destabilizing enzymes. Cell 96:69–78. doi:10.1016/s0092-8674(00)80960-5

14. Gallegos ME, Bargmann CI. 2004. Mechanosensory neurite termination and tiling depend on SAX-2 and the SAX-1 kinase. Neuron 44:239–249. doi:10.1016/j.neuron.2004.09.021

15. Goris A, Boonen S, D’hooghe M-B, Dubois B. 2010. Replication of KIF21B as a susceptibility locus for multiple sclerosis. J Med Genet 47:775–776. doi:10.1136/jmg.2009.075911

16. Guedes-Dias P, Holzbaur ELF. 2019. Axonal transport: Driving synaptic function. Science 366:eaaw9997. doi:10.1126/science.aaw9997

17. Gupta ML, Carvalho P, Roof DM, Pellman D. 2006. Plus end-specific depolymerase activity of Kip3, a kinesin-8 protein, explains its role in positioning the yeast mitotic spindle. Nat Cell Biol 8:913–923. doi:10.1038/ncb1457

18. He M, Subramanian R, Bangs F, Omelchenko T, Liem KF, Kapoor TM, Anderson KV. 2014. The kinesin-4 protein Kif7 regulates mammalian Hedgehog signalling by organizing the cilium tip compartment. Nat Cell Biol 16:663–672. doi:10.1038/ncb2988

19. Hirata K, Yamashita K, Ueno G, Kawano Y, Hasegawa K, Kumasaka T, Yamamoto M. 2019. ZOO: an automatic data-collection system for high-throughput structure analysis in protein microcrystallography. Acta Crystallogr D Struct Biol 75:138–150. doi:10.1107/S2059798318017795

20. Hirokawa N, Nitta R, Okada Y. 2009a. The mechanisms of kinesin motor motility: lessons from the monomeric motor KIF1A. Nat Rev Mol Cell Biol 10:877–884. doi:10.1038/nrm2807

21. Hirokawa N, Noda Y, Tanaka Y, Niwa S. 2009b. Kinesin superfamily motor proteins and intracellular transport. Nat Rev Mol Cell Biol 10:682–696. doi:10.1038/nrm2774

22. Hunter AW, Caplow M, Coy DL, Hancock WO, Diez S, Wordeman L, Howard J. 2003. The kinesin-related protein MCAK is a microtubule depolymerase that forms an ATP-hydrolyzing complex at microtubule ends. Mol Cell 11:445–457. doi:10.1016/s1097-2765(03)00049-2

23. Kabsch W. 2010a. XDS. Acta Crystallogr D Biol Crystallogr 66:125–132. doi:10.1107/S0907444909047337

24. Kabsch W. 2010b. Integration, scaling, space-group assignment and post-refinement. Acta Crystallogr D Biol Crystallogr 66:133–144. doi:10.1107/S0907444909047374

25. Kreft KL, van Meurs M, Wierenga-Wolf AF, Melief M-J, van Strien ME, Hol EM, Oostra BA, Laman JD, Hintzen RQ. 2014. Abundant kif21b is associated with accelerated progression in neurodegenerative diseases. Acta Neuropathol Commun 2:144. doi:10.1186/s40478-014-0144-4

26. Liebschner D, Afonine PV, Moriarty NW, Poon BK, Sobolev OV, Terwilliger TC, Adams PD. 2017. Polder maps: improving OMIT maps by excluding bulk solvent. Acta Crystallogr D Struct Biol 73:148–157. doi:10.1107/S2059798316018210

27. McCoy AJ, Grosse-Kunstleve RW, Adams PD, Winn MD, Storoni LC, Read RJ. 2007. Phaser crystallographic software. J Appl Crystallogr 40:658–674. doi:10.1107/S0021889807021206

28. Mello CC, Kramer JM, Stinchcomb D, Ambros V. 1991. Efficient gene transfer in C.elegans: extrachromosomal maintenance and integration of transforming sequences. The EMBO Journal 10:3959–3970. doi:10.1002/j.1460-2075.1991.tb04966.x

29. Muhia M, Thies E, Labonté D, Ghiretti AE, Gromova KV, Xompero F, Lappe-Siefke C, Hermans-Borgmeyer I, Kuhl D, Schweizer M, Ohana O, Schwarz JR, Holzbaur ELF, Kneussel M. 2016. The Kinesin KIF21B Regulates Microtubule Dynamics and Is Essential for Neuronal Morphology, Synapse Function, and Learning and Memory. Cell Rep 15:968–977. doi:10.1016/j.celrep.2016.03.086

30. Muto E, Sakai H, Kaseda K. 2005. Long-range cooperative binding of kinesin to a microtubule in the presence of ATP. J Cell Biol 168:691–696. doi:10.1083/jcb.200409035

31. Niwa S. 2017. Immobilization of Caenorhabditis elegans to Analyze Intracellular Transport in Neurons. J Vis Exp. doi:10.3791/56690

32. Niwa S, Nakajima K, Miki H, Minato Y, Wang D, Hirokawa N. 2012. KIF19A is a microtubule-depolymerizing kinesin for ciliary length control. Dev Cell 23:1167–1175. doi:10.1016/j.devcel.2012.10.016

33. Ogawa T, Nitta R, Okada Y, Hirokawa N. 2004. A common mechanism for microtubule destabilizers-M type kinesins stabilize curling of the protofilament using the class-specific neck and loops. Cell 116:591–602. doi:10.1016/s0092-8674(04)00129-1

34. O’Hagan R, Chalfie M, Goodman MB. 2005. The MEC-4 DEG/ENaC channel of Caenorhabditis elegans touch receptor neurons transduces mechanical signals. Nat Neurosci 8:43–50. doi:10.1038/nn1362

35. Peet DR, Burroughs NJ, Cross RA. 2018. Kinesin expands and stabilizes the GDP-microtubule lattice. Nat Nanotechnol 13:386–391. doi:10.1038/s41565-018-0084-4

36. Pettersen EF, Goddard TD, Huang CC, Couch GS, Greenblatt DM, Meng EC, Ferrin TE. 2004. UCSF Chimera--a visualization system for exploratory research and analysis. J Comput Chem 25:1605–1612. doi:10.1002/jcc.20084

37. Shima T, Morikawa M, Kaneshiro J, Kambara T, Kamimura S, Yagi T, Iwamoto H, Uemura S, Shigematsu H, Shirouzu M, Ichimura T, Watanabe TM, Nitta R, Okada Y, Hirokawa N. 2018. Kinesin-binding-triggered conformation switching of microtubules contributes to polarized transport. J Cell Biol 217:4164–4183. doi:10.1083/jcb.201711178

38. Tomishige M, Klopfenstein DR, Vale RD. 2002. Conversion of Unc104/KIF1A kinesin into a processive motor after dimerization. Science 297:2263–2267. doi:10.1126/science.1073386

39. Trofimova D, Paydar M, Zara A, Talje L, Kwok BH, Allingham JS. 2018. Ternary complex of Kif2A-bound tandem tubulin heterodimers represents a kinesin-13-mediated microtubule depolymerization reaction intermediate. Nat Commun 9:2628. doi:10.1038/s41467-018-05025-7

40. van der Vaart B, van Riel WE, Doodhi H, Kevenaar JT, Katrukha EA, Gumy L, Bouchet BP, Grigoriev I, Spangler SA, Yu KL, Wulf PS, Wu J, Lansbergen G, van Battum EY, Pasterkamp RJ, Mimori-Kiyosue Y, Demmers J, Olieric N, Maly IV, Hoogenraad CC, Akhmanova A. 2013. CFEOM1-associated kinesin KIF21A is a cortical microtubule growth inhibitor. Dev Cell 27:145–160. doi:10.1016/j.devcel.2013.09.010

41. van Riel WE, Rai A, Bianchi S, Katrukha EA, Liu Q, Heck AJ, Hoogenraad CC, Steinmetz MO, Kapitein LC, Akhmanova A. 2017. Kinesin-4 KIF21B is a potent microtubule pausing factor. Elife 6. doi:10.7554/eLife.24746

42. Varga V, Helenius J, Tanaka K, Hyman AA, Tanaka TU, Howard J. 2006. Yeast kinesin-8 depolymerizes microtubules in a length-dependent manner. Nat Cell Biol 8:957–962. doi:10.1038/ncb1462

43. Wang D, Nitta R, Morikawa M, Yajima H, Inoue S, Shigematsu H, Kikkawa M, Hirokawa N. 2016. Motility and microtubule depolymerization mechanisms of the Kinesin-8 motor, KIF19A. Elife 5. doi:10.7554/eLife.18101

44. Wang W, Cantos-Fernandes S, Lv Y, Kuerban H, Ahmad S, Wang C, Gigant B. 2017. Insight into microtubule disassembly by kinesin-13s from the structure of Kif2C bound to tubulin. Nat Commun 8:70. doi:10.1038/s41467-017-00091-9

45. Waterhouse A, Bertoni M, Bienert S, Studer G, Tauriello G, Gumienny R, Heer FT, de Beer TAP, Rempfer C, Bordoli L, Lepore R, Schwede T. 2018. SWISS-MODEL: homology modelling of protein structures and complexes. Nucleic Acids Res 46:W296– W303. doi:10.1093/nar/gky427

46. Yamada K, Andrews C, Chan W-M, McKeown CA, Magli A, de Berardinis T, Loewenstein A, Lazar M, O’Keefe M, Letson R, London A, Ruttum M, Matsumoto N, Saito N, Morris L, Del Monte M, Johnson RH, Uyama E, Houtman WA, de Vries B, Carlow TJ, Hart BL, Krawiecki N, Shoffner J, Vogel MC, Katowitz J, Goldstein SM, Levin AV, Sener EC, Ozturk BT, Akarsu AN, Brodsky MC, Hanisch F, Cruse RP, Zubcov AA, Robb RM, Roggenkäemper P, Gottlob I, Kowal L, Battu R, Traboulsi EI, Franceschini P, Newlin A, Demer JL, Engle EC. 2003. Heterozygous mutations of the kinesin KIF21A in congenital fibrosis of the extraocular muscles type 1 (CFEOM1). Nat Genet 35:318–321. doi:10.1038/ng1261

47. Yamashita K, Hirata K, Yamamoto M. 2018. KAMO: towards automated data processing for microcrystals. Acta Crystallogr D Struct Biol 74:441–449. doi:10.1107/S2059798318004576

48. Yue Y, Blasius TL, Zhang S, Jariwala S, Walker B, Grant BJ, Cochran JC, Verhey KJ. 2018. Altered chemomechanical coupling causes impaired motility of the kinesin-4 motors KIF27 and KIF7. J Cell Biol 217:1319–1334. doi:10.1083/jcb.201708179

